# *Inseparable/IER3IP1* are essential for cytokinesis in *Drosophila* neuroblast and human cells

**DOI:** 10.1101/2024.07.20.604396

**Authors:** Aishwarya Arun Kakade, Sachin Gupta, Reshmi Varghese, Harikrishna Adicherla, Sonal Nagarkar-Jaiswal

## Abstract

To unveil the molecular players that maintain neural stem cell homeostasis, we conducted a genetic screen in *Drosophila* and isolated an uncharacterized gene that we named *Inseparable* (*Insep*). *Insep* is the *Drosophila* homologue of human *IER3IP1*, a gene associated with Microcephaly, Epilepsy, and Neonatal Diabetes Syndrome (MEDS-1). We show that *Insep* loss leads to early larval lethality with small brains and these phenotypes can be rescued by expressing IER3IP1 indicating that their biological function is conserved through evolution. The *Insep* deficient neuroblasts fail to complete cytokinesis and show excessive accumulation of Rab11 vesicles in the cytoplasm. Similarly, *IER3IP1* depletion in human cells leads to cytokinesis failure and accumulation of Rab11 vesicles. Insep and IER3IP1 localize to Rab11 vesicles and interact with Rab11. The pathogenic mutations in IER3IP1 perturb its localization to Rab11 vesicles and interaction with Rab11. These results suggest that Insep and IER3IP1 work along with Rab11 and may regulate fusion of Rab11 vesicles to the advancing furrow during cytokinesis.

## Introduction

Neural stem cells (NSCs), self-renew to maintain their pool and differentiate into functionally and morphologically distinct neurons and glia during brain development (1). The proper brain development depends on a precise balance between the amplification of NSCs and the constant generation of differentiated progeny. De-deregulation in NSC maintenance can lead to conditions such as microcephaly or brain tumour (2).

*Drosophila* neural stem cells, neuroblasts (NBs) have emerged as an unparalleled *in vivo* platform to study NSC self-renewal and differentiation during development (1, 3). The fly brain lobe contains type I, type II, mushroom body, and optic lobe NBs; among these, the type I and II neuroblasts generate the majority of larval and adult neurons and glia (1). The Type I NBs undergo asymmetric division and give rise to a daughter neuroblast and a ganglion mother cell (GMC), which eventually divides symmetrically and gives rise to two neurons or two glial cells (1, 3). The type II NBs divide asymmetrically and produce one daughter NB and an intermediate neural precursor (INP). INP further undergoes a few rounds of asymmetric divisions, giving rise to one INP and one GMC (1, 3). The continuous asymmetric division maintains the NB pool and generates neurons and glia temporally.

A crucial step for NB asymmetric division is the establishment of apical/basal polarity (4). In interphase, apical polarity complex proteins and cell fate determinants are distributed in cytoplasm (5, 6). As NBs enter prophase, the apical polarity complex proteins Par3/ Bazooka, aPKC, and Par-6 start to localize at the apical cortex, which drives the localization of the cell-fate determinants and their adaptor proteins to the basal cortex (5, 7–14). During division, the apical polarity proteins and cell fate determinants segregate to the daughter NB and GMC, respectively (1, 3). In GMC, Mira is degraded and Pros translocates to the nucleus where it suppresses cell cycle genes and induces the expression of neuronal differentiation genes (15). Loss of aPKC causes mislocalization of Mira to the entire cell cortex resulting in segregation of Mira and Pros to both the daughter cells leading to the premature differentiation of NB (8). Similarly, Mira loss of function mutants fail to anchor Pros at the basal cortex, as a result Pros accumulate in the cytoplasm, and upon NB division, Pros is distributed to both NB and GMC, causing their premature differentiation into neurons leading to the exhaustion of NBs and small brain phenotype (13).

In humans, reduced proliferation and premature differentiation of neural progenitor cells are known to cause microcephaly, a clinical condition that is categorized as neurodevelopmental impairment when a child’s head circumference falls 1.96 standard deviations (SDs) or more below the average for their age and sex (16–22). One such condition is MEDS1 (Microcephaly, epilepsy, and diabetes syndrome-1, OMIM# 614231) which is an autosomal recessive inherited disorder. MEDS-1 is characterized by microcephaly with a reduced number of gyri, neonatal epilepsy, and infantile diabetes and has been shown to be caused by mutations in *IER3IP1* (23–27)

IER3IP1 is a small protein with two transmembrane domains and was shown to localize to ER in human cells (28). The yeast homolog of IER3IP1, YOS1P, localizes to ER, Golgi, and COPII vesicles, and its depletion blocks early secretory pathway i.e. ER to Golgi transport (29). In mice, IER3IP1 is expressed in pancreatic beta cells and its loss leads to proinsulin misfolding, ER stress, reduced proliferation of beta cells and apoptosis (23, 30, 31). A recent study showed that human brain organoids derived from IER3IP1 deficient iPSCs are smaller than controls, they show abnormal neural progenitor localization and defects in extracellular matrix protein secretion and tissue integrity (32). However, the precise pathogenic mechanism underlying MEDS-1 due to the loss of IER3P1 is not clear.

In this study, we characterize the fly orthologue of *IER3IP1*, which we named *Inseparable* (*Insep*). We show that *Insep* is required for brain development in flies and the function of *Insep* and *IER3IP1* is conserved. We found that *Insep* and *IER3IP1* are required for the completion of cytokinesis. We discovered that Insep and IER3IP1 interact with Rab11, a GTPase that has been shown to be required for fusion of vesicles to the advancing furrow during cytokinesis. The disease-causing mutations (V21G and L78P) perturb the localization of IER3IP1 to Rab11 vesicles and interaction with Rab11. Our results suggest that Insep and IER3IP1 work along with Rab11 in completing cytokinesis, possibly by regulating the fusion of Rab11 vesicles to the advancing furrow during cytokinesis.

## Results

### *Insep* is required for brain development and survival in flies

We conducted a gene expression-based screen using CRIMIC based *T2A-GAL4* gene trap driver lines (33) and *UAS-EGFP* reporter to identify genes that are involved in brain development in *Drosophila.* Through this screen, we identified *CG32069*, an uncharacterized gene that is expressed in larval brains (Figure S1 A and B). We named it *Inseparable* (*Insep*) based on the phenotype described later (Figure 2-A and B). To explore the role of *Insep* in brain development, we adapted three strategies to create loss-of-function mutants. First, we generated a null allele of *Insep* (*Insep^N^*) in which we deleted the entire coding region and replaced it with a 3XP3-EGFP cassette by CRISPR/Cas9 mediated homologous recombination (Figure 1-A). Second, we performed conditional knockout of *Insep* by expressing Cas9 under Actin promoter (*Act-Cas9*) in larvae that carry a transgene to express guide RNA targeting *Insep* under U6 promoter, from hereon we will call it *Insep CN*. Third, we utilized an RNAi line (v9202) to knock down *Insep* in developing brains using *Inscutable-GAL4* driver (*Insep RI*). We observed a developmental delay and larval lethality in all three conditions (Figure 1- B).

**Figure 1:**
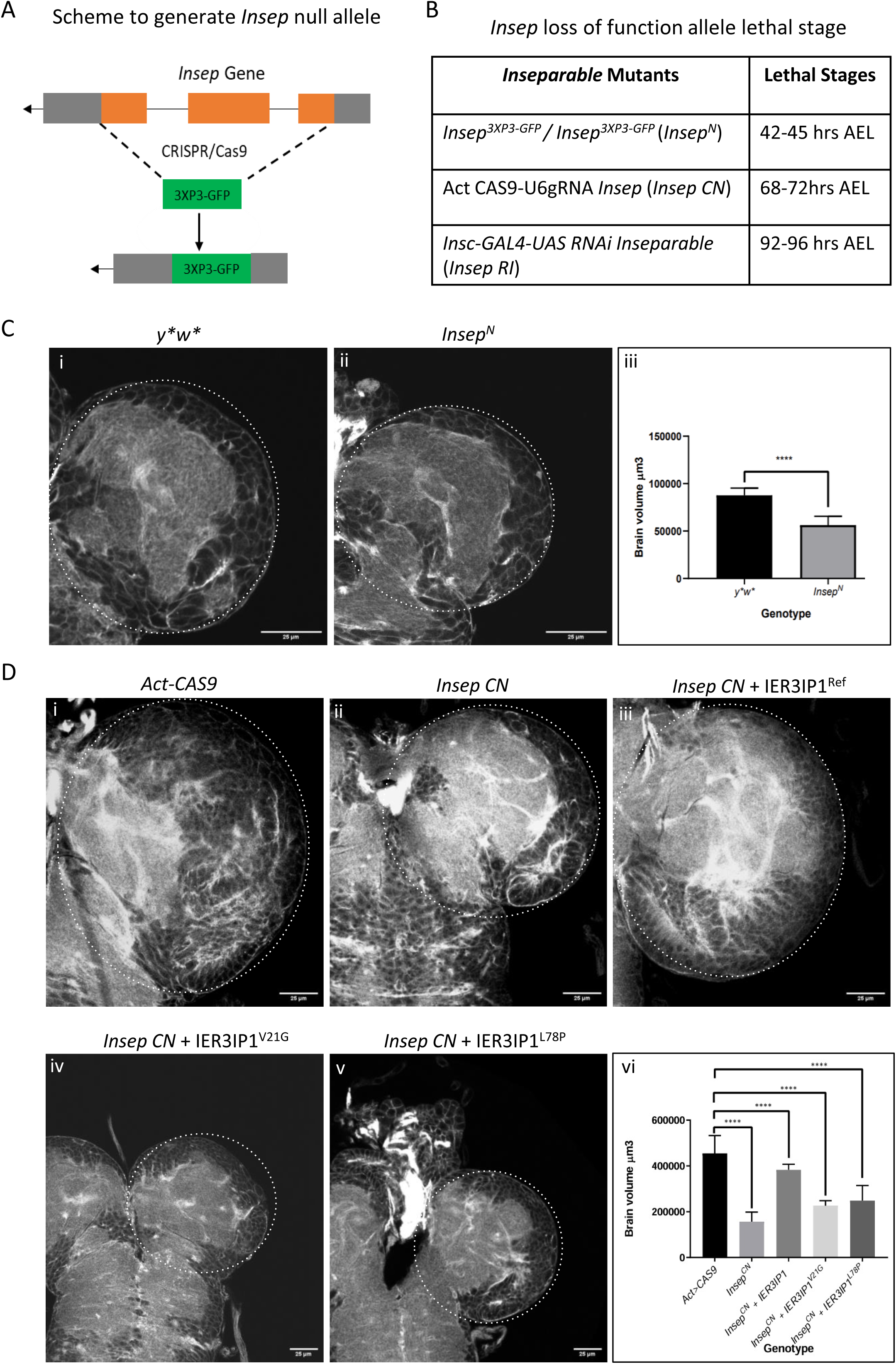
*Inseparable* loss leads to larval lethality with small brains. A. Scheme depicting generation of *Inseparable* null mutant using *CRISPR/Cas9* system, where *Insep* coding region was replaced by a 3XP3-GFP cassette B. Lethal stages of *Insep* loss of function mutants*; Insep^N^* -null alelle*, Insep CN* -conditional null and *Insep RI -*RNAi mediated knock down C. Brain lobe size of (i) *y*w\** control (ii) *Insep^N^*, larval brains are stained with Phalloidin (iii) quantification of (i, n=17) and (ii, n=16), ∗p<0.01, ∗∗∗p<0.001, ∗∗∗∗p<0.0001 (i) *Act-Cas9* control (ii) *Insep CN* (iii) *Insep CN* larvae expressing Reference IER3IP1, IER3IP1^Ref^ (iv) *Insep CN* larvae expressing V21G variant of IER3IP1, IER3IP1^V21G^ (v) *Insep CN* larvae expressing L78P variant of IER3IP1, IER3IP1^L78P^, larval brains are stained with Phalloidin. (vi) Quantification of (i, n=15), (ii, n=15), (iii, n=10), (iv, n=8) and (v, n=8), ∗∗p<0.01, ∗∗∗p<0.001, ∗∗∗∗p<0.0001. n=number of brains

**Figure 2:**
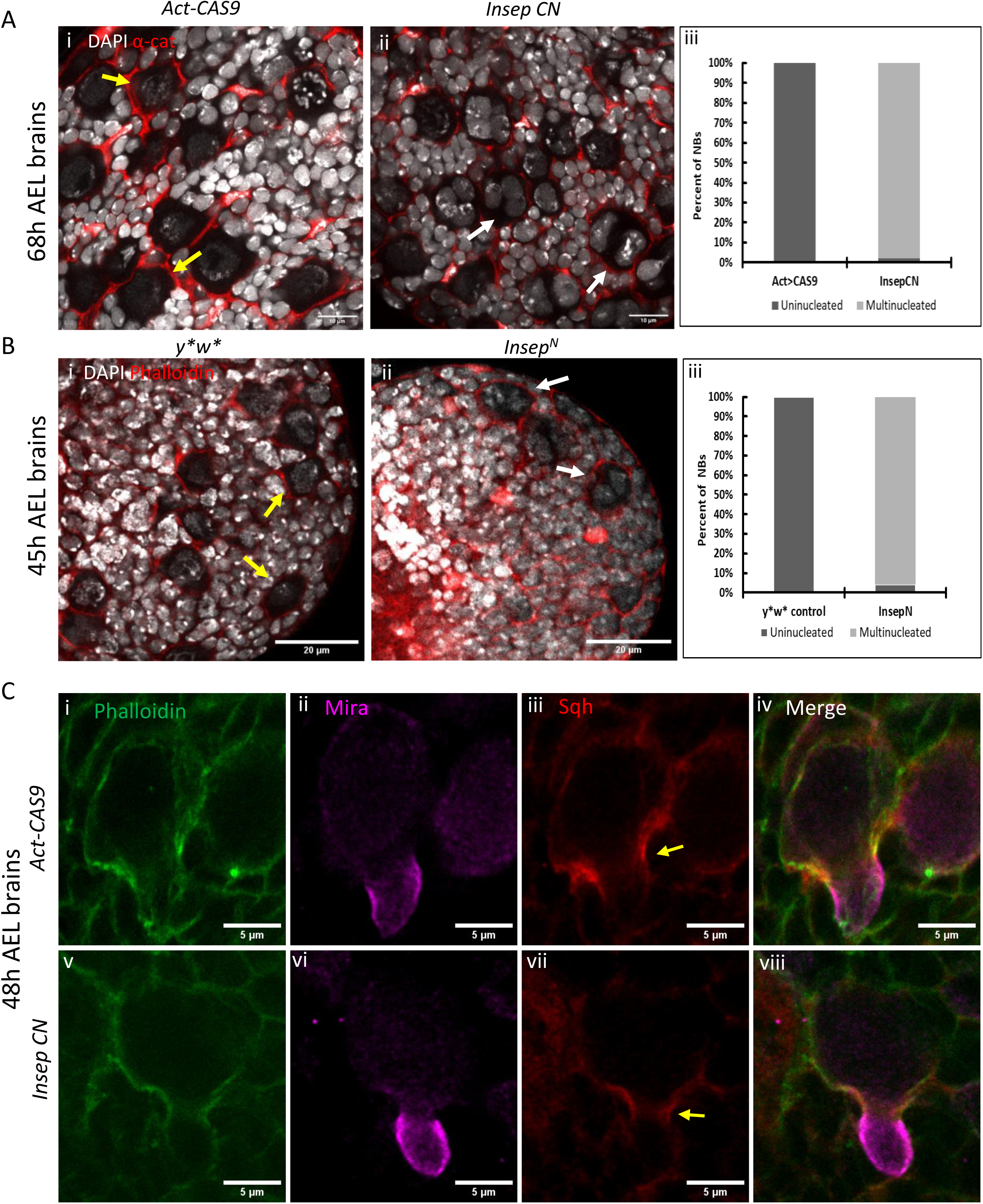
*Insep* mutant NBs fail to complete cytokinesis A. Confocal images of brain of 68h AEL larvae from *Act-Cas9* controls (1, n=12) and *Insep CN* (ii, n=14) stained with DAPI (gray) and α-catenin (red). Yellow arrows indicate NB with single nucleus (i), white arrows indicate multinucleated NBs (ii), (iii) shows quantification of (i and ii). Scale bar=10µm B. Confocal images of brain of 45h AEL larvae from *y*w\** controls (i, n=12) and *Insep^N^* (ii, n=14) stained with DAPI (gray) and Phalloidin (red). Yellow arrows indicate NB with single nucleus (i), white arrows indicate multinucleated NBs (ii), (iii) shows quantification for (i and ii) .Scale bar=20µm C. Confocal images of brain from *Act-Cas9* controls (i, ii, iii, iv, n=3) and *Insep CN* (v, vi, vii, viii, n=5) 48h AEL larvae stained with Phalloidin (green, i, iv, v, viii), Mira (magenta, ii, iv, vi and viii) and Sqh (red iii, iv, vii and viii), yellow arrows indicate Sqh localization at the cleavage furrow. Scale bar=5µm. n = number of brains

Further, we analyzed the brain sizes of *Insep^N^* and *Insep CN* animals. We found that the brains of these animals are significantly smaller than the respective controls (Figure 1-C, i-iii, and Figure 1-D, i, ii and vi). To further confirm that small brains and larval lethality are due to the loss of *Insep*, we generated transgenic lines expressing HA::Insep protein (Figure S1-C, i) and performed rescue experiments. We expressed HA::Insep in *Insep^N^*and *Insep CN* animals ubiquitously using the *Tubulin-GAL4* driver (*Tub-GAL4>UAS-HA::Insep*). We found that expression of HA::Insep rescues the small brain phenotype and larval lethality indicating that these phenotypes are due to the loss of *Insep* and HA::Insep is a functional protein.

### *Insep* shows functional conservation with human *IER3IP1*

*Insep* is a predicted orthologue of *IER3IP1* (53% identity, 73% similarity) and mutations in *IER3IP1* in humans are linked to microcephaly (24–27), which is an analogous phenotype caused by the loss of *Insep* in flies suggesting a functional conservation between *IER3IP1* and *Insep*. To confirm the functional conservation, we decided to check the rescue of *Insep* loss of function phenotype by expressing IER3IP1. Thus we generated transgenic lines to express human reference *IER3IP1* cDNA (*IER3IP1^Ref^*) and disease associated variants reported in MEDS-1, *IER3IP1^V21G^* and *IER3IP1^L78^* (23, 24, 26, 27), using *UAS-GAL4* system which allows tissue specific expression (Figure S1-C, ii, iii and iv). We expressed IER3IP1 in *Insep CN* and *Insep^N^* animals ubiquitously using *Tubulin-GAL4 driver* (*Tub-GAL4>UAS-IER3IP1^Ref^* or *^V21G^* or *^L78P^*). We found that *IER3IP1^Ref^* rescues the larval lethality associated with *Insep* loss, whereas, the disease variants *IER3IP1^V21G^* and *IER3IP1^L78P^* do not rescue the lethality (Figure 1-D, iii, iv, v and vi). We also observed that levels of IER3IP1^V21G^ and IER3IP1^L78P^ proteins were much lower than IER3IP1^Ref^ suggesting that these mutations may affect IER3IP1 protein stability (Figure S1-C ii, iii and iv). These results confirm that *Insep-IER3IP1* function is evolutionarily conserved and the disease associated variants are severe loss of function alleles.

### *Insep* mutant NBs fail to complete cytokinesis

Neuroblasts undergo multiple rounds of asymmetric division and generate neurons and glia; this timely division of NBs is crucial for brain development (1–3). We checked whether *Insep* loss affects NBs division and found that mutant NBs are multinucleated. Upon detailed analysis, we found in *Insep CN* about 94% of mutant NBs in 48h AEL and about 98% of mutant NBs from 68h AEL larval brains contain more than one nucleus (Figure 2-A, i, ii and iii). Similarly, in 45h AEL *Insep^N^*larval brains more than about 96% of mutant NBs were multinucleated (Figure 2-B, i, ii and iii). This data suggests that *Insep* deficient NBs fail to complete cytokinesis, which may result in early termination of neurogenesis leading to small brains.

Neuroblasts establish apico/basal polarity during asymmetric division that ensures self-renewal and differentiation. We sought to check whether loss of *Insep* affects NBs polarity (Figure S2-A and B). We found about 6% NBs of 48h AEL larval brains had apically localized aPKC and basally localized Mira and Pros and 93% had aPKC, Mira, and Pros mislocalized to the cytoplasm and remaining NBs may be in interphase. In 68h AEL larval brains, 1% NBs had proper localization of aPKC and Mira, 98% NBs showed mislocalization to the cytoplasm and remaining NBs may be in interphase (Figure S2-A v, vi, vii and viii). In *Insep^N^*, 96% NBs of 45h AEL larval brains had aPKC, Mira, and Pros mislocalized to the cytoplasm (Figure S2-B v, vi, vii and viii). Since, we find NBs with aPKC/Mira localization even at 68h AEL, a stage after which larvae die, this suggests that mutant NBs establish polarity and initiate asymmetric division, however, they fail to complete cytokinesis. The cytokinesis failure in NBs results in redistribution of aPKC and Mira in cytoplasm, where, in some NBs Mira is lost and Pros translocates to the nucleus driving expression of neural genes that leads to reduction in NB pool (Figure S2-A and B, vi and vii)). Based on these results we conclude that microcephaly-like phenotype in flies is caused by failure to complete cytokinesis and a reduction in NB pool is due to *Insep* loss.

### *Insep* is not required for actomyosin assembly

One of the critical steps that precede cytokinesis is the assembly of a contractile actomyosin ring, which then undergoes constriction that leads to the ingression of the cleavage furrow that eventually divides the cell into two daughter cells (34–37). Therefore, we checked whether the cytokinesis failure in *Insep* mutants NBs is due to a defect in actomyosin ring assembly and/or constriction. We examined Spaghetti squash (Sqh), the regulatory light chain of non-muscle Myosin II in *Drosophila* (11, 12), to observe actomyosin ring assembly and constriction in NBs. We generated fly stocks expressing *Sqh-mCherry* in *Insep CN* background and performed immunostaining on brains from 48h AEL larvae, in which we find around 6% NBs initiating asymmetric division. We found that similar to control NBs (Figure 2-C, i-iv), the mutant NBs assemble actomyosin ring and initiate constriction, however, eventually, they regress and fail to complete cytokinesis (Figure 2-C, v-viii). These results show that *Insep* may not be required for actomyosin assembly in NBs.

### *Insep* mutant NBs accumulate Rab11 vesicles

Membrane trafficking is crucial for cytokinesis, secretory and recycling vesicles move towards and fuse to the ingressing furrow to deliver new membrane and proteins required for the progression of the cleavage furrow and subsequent abscission (38–45). *Insep* orthologues *IER3IP1* and *Yos1P* have also been associated with vesicle trafficking (38, 40). Therefore, we investigated vesicle trafficking in *Insep* mutant NBs. We examined Rab11, a small GTPase, which is implicated in both recycling of endocytic vesicle and secretory pathways, and has been shown to be required for cytokinesis completion (38, 40). We found that compared to the control NBs where we occasionally see Rab11 puncta in the cytoplasm (Figure 3-A, i-iv and Figure S3-A, i-iv), *Insep CN* and *Insep^N^* NBs have increased levels of Rab11 in the cytoplasm with a large blob-like structure (Figure 3-A, v-viii and Figure S3-A, vi-viii). We investigated whether Rab11 is accumulated in ER. We used KDEL to mark ER and did not find KDEL signal localising to the Rab11 blobs suggesting Rab11 is not accumulated in ER (Figure S3-B).

**Figure 3:**
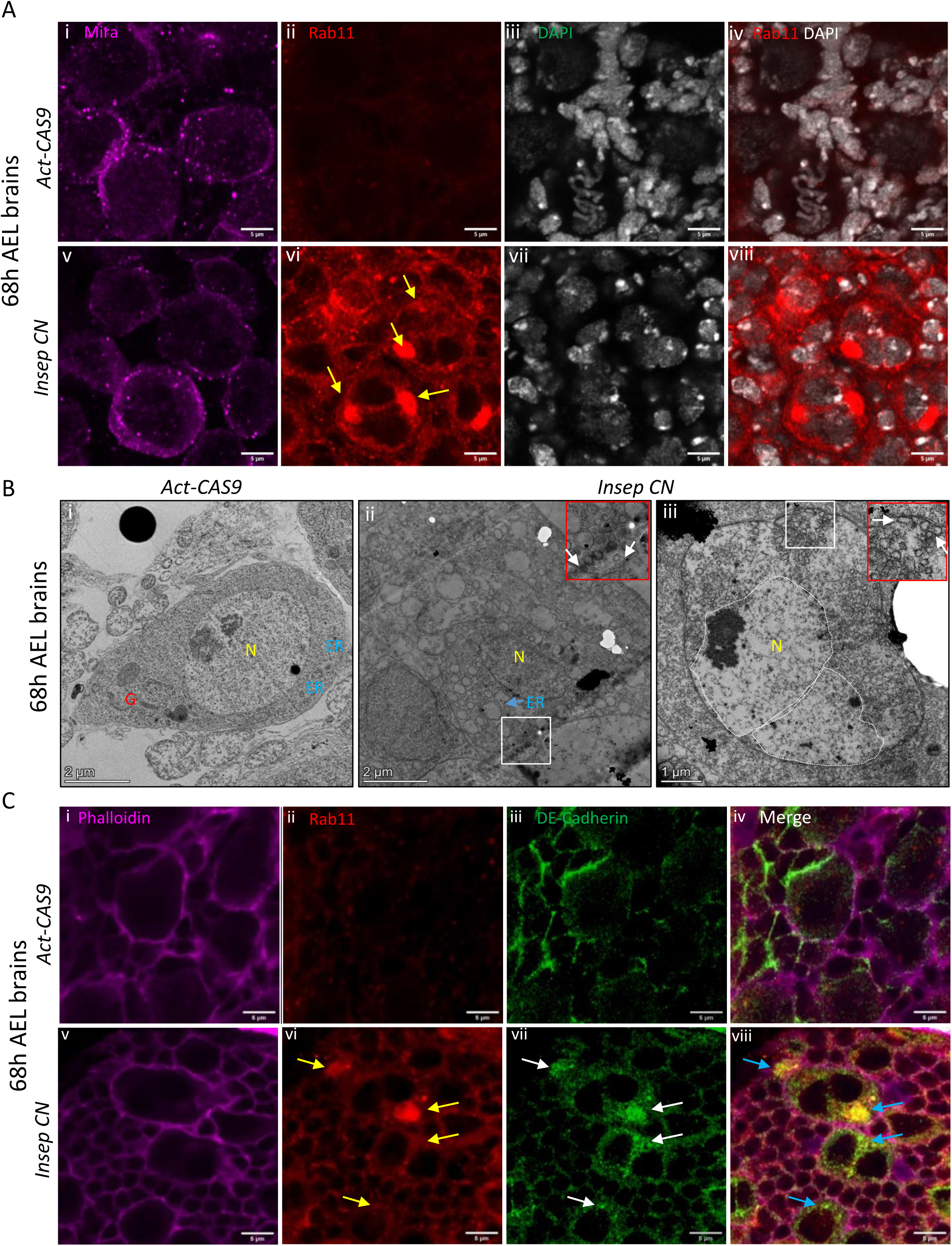
*Insep* mutant NBs accumulate Rab11 vesicles. A. Confocal images of 68h AEL brain from *Act-Cas9* controls (i-iv, n=11) and *Insep CN* (v-viii, n=17) stained with Mira (magenta, I and v), Rab11 (red, ii, iv, vi, viii) and DAPI (gray, iii, iv, vii, viii), yellow arrows show accumulation of rab11 (vi). Scale bar=5µm. B. Electron microscopy images of control (i, n=8) and *Insep CN* NB (ii, iii, n=8). Blue arrow in (ii) indicate widened ER, white arrows indicate vesicles accumulated near plasma membrane (ii and iii), red insets in show magnified view of white inset in the respective image. N-Nucleus, G-Golgi, ER-Endoplasmic Reticulum, Scale bar = 2µm (i, ii) and 1µm (iii) C. Confocal images of brain from *Act-Cas9* controls (i, ii, iii, iv, n=7) and *Insep CN* (v, vi, vii, viii, n=6) 68h AEL larval brains stained with Phalloidin (magenta, i, iv, v and viii), Rab11 (red, ii, iv, vi and viii), DE-Cad (green, iii, iv, vii and viii), yellow arrows show accumulation of Rab11 vesicle in (vi), white arrows show accumulation of DE-Cad in (vii) and blue arrows in (viii) shows co-localization of Rab11 and DE-Cad. Scale bar=5µm

Next, we performed transmission electron microscopy on *Insep CN* brains. We observed that ER in mutant NBs was widened compared to the controls (Figure 3-B, i and ii), a phenotype that has been reported for *IER3IP1* KO organoids and *Yos1P* mutants (29, 32). We also found that the number of vesicles was drastically increased in mutant NBs, vesicles were present in the cytoplasm and accumulated close to the plasma membrane (Figure 3-B, ii and iii), and in some cells vesicles were present in clusters (Figure 3-B, iii, inset). To check whether the accumulated vesicles are Rab11 vesicles, we analyzed DE-Cadherin, a Rab11 vesicle cargo protein (17,18,19). We observed an increase in DE-Cadherin levels in *Insep* mutant NBs (Figure 3-C and Figure S3-C), and found that DE-Cadherin co-localizes to Rab11 blobs suggesting that these are indeed clusters of Rab11 vesicles (Figure 3-C, vi-viii).

We checked whether *Insep* loss affects Rab11 vesicle transport to the ingressing furrow (Figure S5), We performed immunostaining for Rab11 on brains from 48n AEL *Insep CN* larvae in which few NBs have initiated asymmetric division. We found that the mutant NBs accumulate Rab11 vesicles at the ingressing cleavage furrow and mid body similar to the control NBs (Figure S4). Our data suggests that *Insep* loss may not affect Rab11 vesicle formation and transport to the ingressing furrow, and the cytokinesis failure could be due to defects in the vesicle tethering or fusion.

### *IER3IP1* is required for cytokinesis in human cells

*IER3IP^Ref^* rescues the brain defects and larval lethality associated with *Insep* mutants, indicating that *Insep’*s function is conserved in humans. Therefore, we checked whether *IER3IP1* loss also leads to cytokinesis defects in human cells. We designed shRNA constructs to knock-down (KD) *IER3IP1* in HCT116 and HEK293 cells; we observed efficient knock-down in both cell lines. (Figure S5-A, i and ii). We found that in unsynchronised *IER3IP1* KD HCT116 cells about 28% of cells have cytokinesis defects (n= 254, 2 experiments) and control cells had 3% cells with cytokinesis defect (n=265, 2 experiments). However, when we treat HCT116 cells with nocodazole to enrich the cells with cytokinesis defect, we found about 48% of IER3IP1 KD cells had cytokinesis failure (n=357, 3 experiments) and control cells had 2% cells with cytokinesis defect (n=373, 3 experiments) (Figure 4-A). Similarly, in unsynchronised HEK293 cells 31% of cells have cytokinesis failure (n=258, 3 experiments) and control cells had 2% cells with cytokinesis failure (n=257, 3 experiments). These results indicate that *IER3IP1* is required for completion of cytokinesis in human cells.

**Figure 4:**
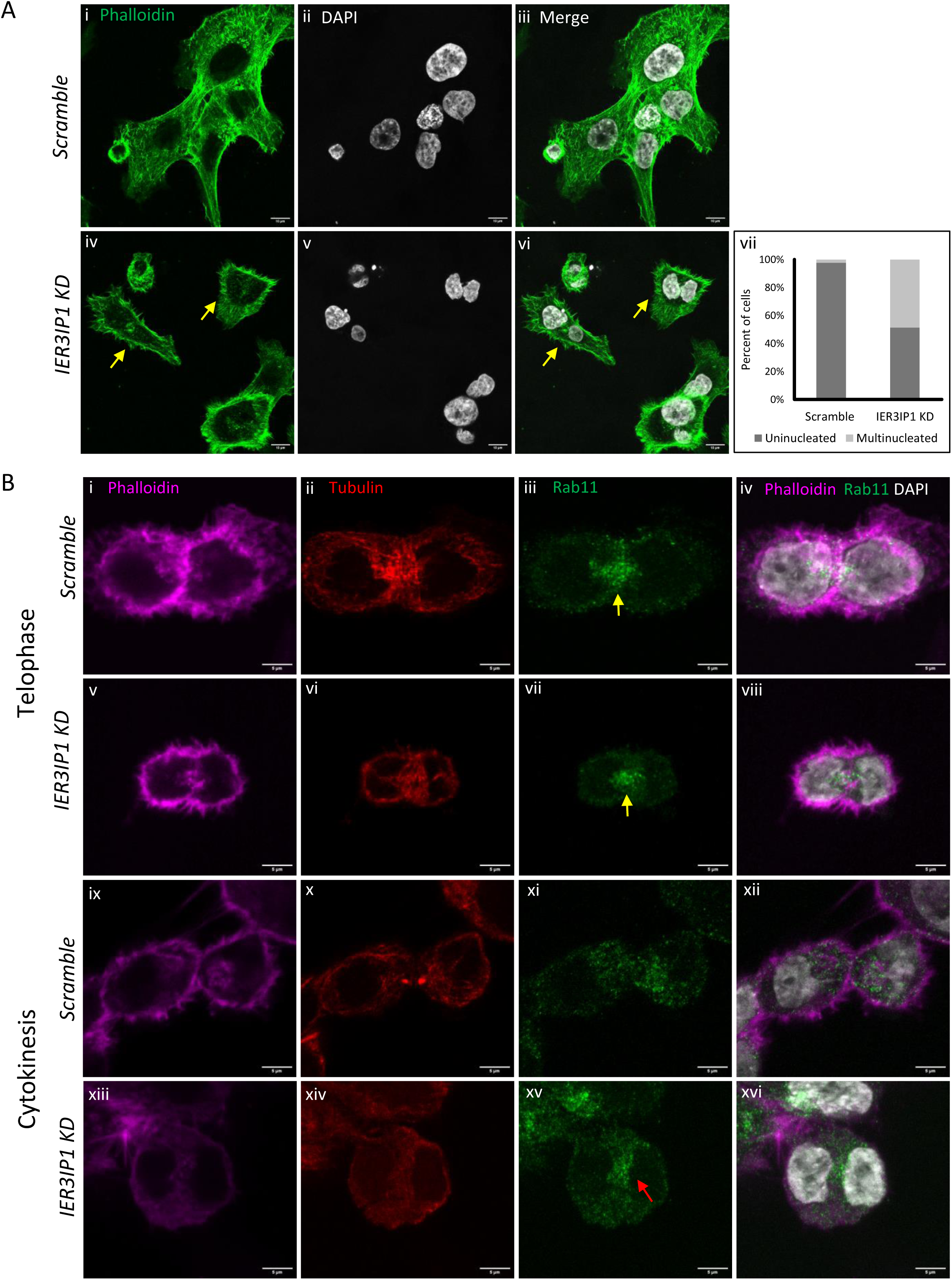
Loss of *IER3IP1* leads to cytokinesis failure in HCT116 cells. A. Confocal images of scramble control (i-iii) and *IER3IP1* shRNA treated HCT116 cells (iv-vi) stained with Phalloidin and DAPI. Yellow arrows in iv and vi indicate cells with cytokinesis failure, vii shows quantification of (i-vi). Scale bar=10µm. B. Confocal images of scramble control (i-iv and ix-xii) and *IER3IP1* KD (v-viii and xiii-xvi) cells stained with Phalloidin, Tubulin, Rab11 and DAPI. Yellow arrows indicate Rab11 accumulation at the ingressing furrow in scramble control cells (iii and iv) and IER3IP1 KD cells (vii and viii), red arrow indicates accumulation of Rab11 vesicles at the midbody in IER3IP1 KD cells that shows cytokinesis failure. Scale bar=5µm

### *IER3IP1* loss does not affect Rab11 vesicle trafficking to the ingressing furrow

To investigate whether Rab11 vesicle trafficking is affected upon *IER3IP1* loss, we examined Rab11 during different phases of mitosis in HCT116 cells after synchronization with nocodazole. We observed that during anaphase, in both scramble control (Figure S5-C, i-iv, n=60) and *IER3IP1* KD (Figure S5-C, v-viii, n= 71), Rab11 vesicles are distributed throughout the cytoplasm. During telophase, Rab11 vesicles travel towards and accumulate at the ingressing furrow and midbody in both scramble control (Figure 4-B, i-iv, n=58) and *IER3IP1* KD (Figure 4-C, v-viii, n=68). In scramble control, when cells complete cytokinesis Rab11 vesicles are distributed throughout the cytoplasm (Figure 4-B, ix-xii, and xii, n=57). In *IER3IP1* KD cells the cleavage furrow is regressed, cells fail to complete cytokinesis and the Rab11 vesicles remain at the midbody position (Figure 4-B xiii-xvi, n=61). Hence, we conclude that *IER3IP1* is not required for the transport of Rab11 vesicles to the progressing cleavage furrow or midbody, but may be involved in vesicle tethering or fusion.

### Insep and IER3IP1 localize to Rab11 vesicles and interact with Rab11

Our data suggests that Insep/IER3IP1 may be involved in Rab11 vesicle docking/tethering or fusion at the ingressing furrow. Therefore, we investigated whether Insep and IER3IP1 localize to Rab11 vesicles. We expressed HA::Insep in wild-type larvae using the *Actin-GAL4* driver (*Act-GAL4>UAS-HA::Insep*) and performed immunostainings using antibodies against HA tag and Rab11. We found that HA::Insep co-localises with Rab11 (Figure 5-A). To check the colocalization between IER3IP1 and Rab11 in human cells, we generated constructs to express HA::IER3IP1 and performed co-staining for HA and Rab11. We observed HA-IER3IP1 co-localizes with Rab11 (Figure 5-B). Next, we checked whether Insep and IER3IP1 interact with Rab11. We performed co-immunoprecipitation (co-IP) experiments on extracts from *Drosophila* S2 cells expressing HA::Insep (*Act-GAL4>UAS-HA::Insep*) and found that Insep interacts with Rab11 (Figure 5-C, i). We further checked whether this interaction is conserved in humans, for this, we performed co-immunoprecipitation experiments on extracts from HCT116 and HEK293 cells expressing HA::IER3IP1 using anti-HA antibody. We observed that HA::IER3IP1 immunoprecipitated Rab11 (Figure 5-C-ii and Figure S6-A). To confirm this interaction, we performed reciprocal co-IP from lysates from cells expressing either mCherry-Rab11 or HA::Rab11 using anti-mCherry or anti-HA antibodies. We found mCherry::Rab11 and HA::Rab11 immunoprecipitated IER3IP1 (Figure 5-C iii and Figure S6-B). These results confirm that Insep and IER3IP1 interact with Rab11 and suggest that they may be working along with Rab11 in vesicle docking/tethering or fusion.

**Figure 5:**
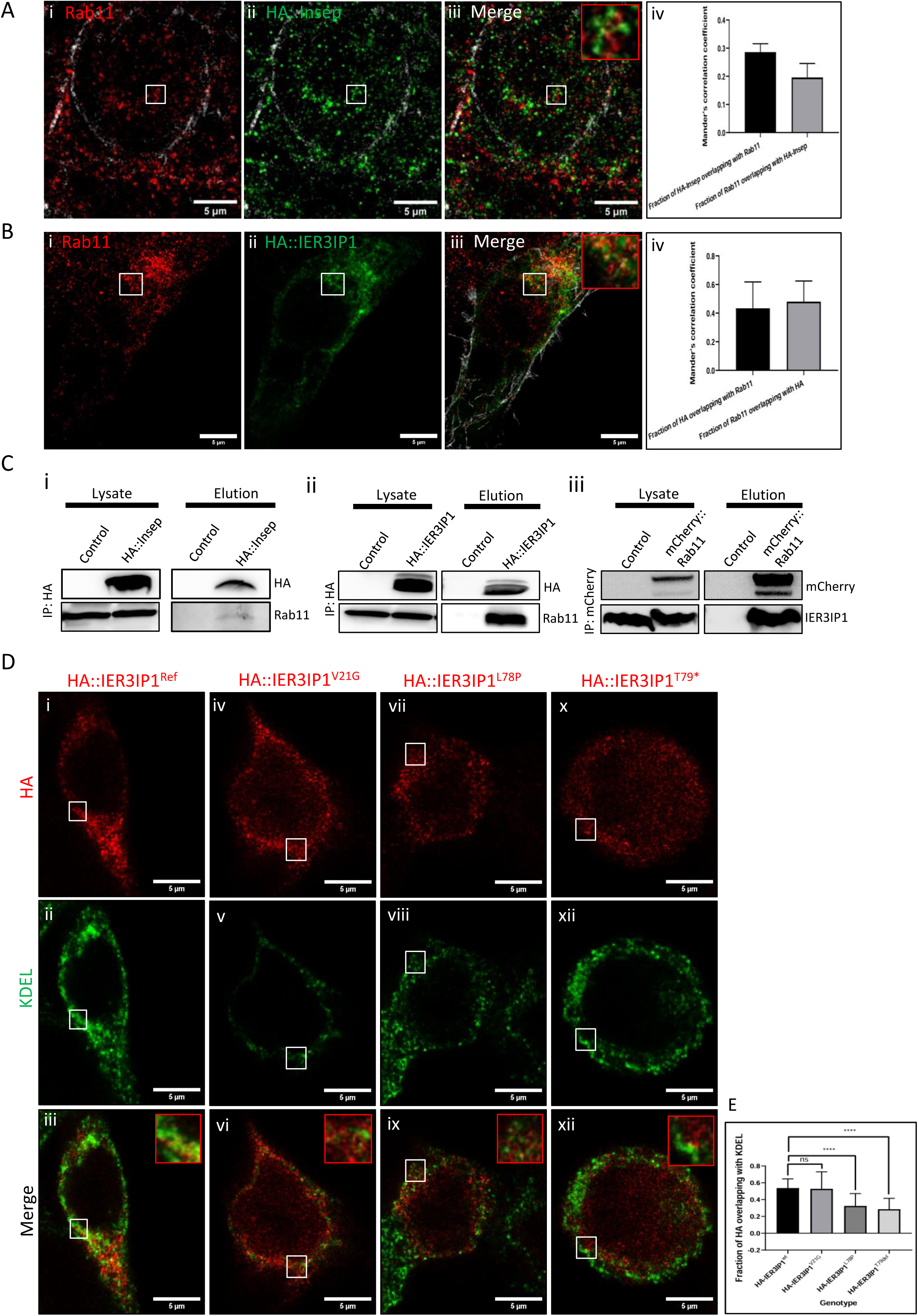
Insep and IER3IP1 localize to Rab11 vesicles and interact with Rab11. A. Brains from L3 larvae expressing HA::Insep were stained with antibodies against Rab11 (red, i and iii) and HA (green, ii and iii), red insets in iii shows the magnified view of the white inset, (iv) shows Mander’s correlation coefficients for HA::Insep overlap with Rab11 and Rab11 overlap with HA::Insep. Scale bar=5µm. B. HCT116 cells expressing HA::IER3IP1 were stained with anti-Rab11 (red, i and iii) and anti-HA (green, ii and iii) antibodies, red inset in (iii) shows the magnified view of the white inset, (iv) shows Mander’s correlation coefficients for HA::IER3IP1 overlap with Rab11 and Rab11 overlap with HA::IER3IP1, scale bar=5µm C. Immunoprecipitation assays showing Rab11 was immunoprecipitated from lysates of S2 cells expressing HA::Insep (i) and HCT116 cells expressing HA::IER3IP1 (ii) using anti-HA beads. Blots were probed with anti-mCherry and anti-IER3IP1 antibodies. iii shows IER3IP1 was immunoprecipitated from lysates of HCT116 cells expressing mCherry::Rab11 using anti-RFP beads and blots were probed with anti-mCherry and anti-IER3IP1 antibodies. D. HCT116 cells expressing *IER3IP1 Ref cDNA* (i, ii and iii) and disease associated variants *V21G* (iv, v and vi), *L78P* (vii, viii and ix) and *T79\** (x, xi and xii) were stained with anti-HA and anti-KDEL antibodies, red insets shows the magnified view of the white inset in the same panel, scale bar=5µm. E. Shows Mander’s correlation coefficient for fraction of HA overlapping with KDEL for HA-IER3IP1^Ref^ , HA-IER3IP1 ^V21G^, HA-IER3IP1 ^L78P^ and HA-IER3IP1 ^T79*.^ ∗∗p<0.01, ∗∗∗p<0.001, ∗∗∗∗p<0.0001.

### Pathogenic mutations perturb localization of IER3IP1 to ER and Rab11 vesicles, and interaction with Rab11

Next, we analysed whether human disease variants affect the localization of IER3IP1 to ER or Rab11 vesicles and their interaction with Rab11. Four different variants in IER3IP1 have been shown to cause MEDS-1 to date. These include two homozygous substitutions, Alanine (A) to Valine (V) at position 18 (A18V) and Valine (V) to Glycine (G) at position 21 (V21G) that falls in N-terminal TM domain, a frameshift allele at nucleotide 79 (T79*), a Leucine (L) to Proline (P) substitution at position 78 that falls in the C-terminal TM domain (23, 24, 26, 27). We expressed *HA::IER3IP1^Ref^*, *HA::IER3IP1^V21G^*, *HA::IER3IP1^L78P^* and *HA::IER3IP1^T79*^* in HCT116 cells. We observed that compared to HA::IER3IP1^Ref^ the levels of HA::IER3IP1^V21G^, and HA::IER3IP1^L78P^ were drastically reduced suggesting that these mutations affect the protein stability (Figure S6-C and D). Further, we checked whether the mutant proteins localized to ER and Rab11 by immunostaining with anti-HA and KDEL or Rab11 antibodies. We found that HA::IER3IP1^Ref^ (Figure 5-D, i-iii and E) and HA::IER3IP1^V21G^ (Figure 5-D, iv-vi and E) localize to ER, HA::IER3IP1^L78P^ show partial localization (Figure 5-D, vii-ix and E), whereas HA::IER3IP1^T79*^ is distributed in the cytoplasm and does not show any localization to ER (Figure 5-D, x-xii and E). Further, we found that HA::IER3IP1^Ref^ colocalizes with Rab11 (Figure 6-A, i-iii and B), HA::IER3IP1^V21G^ (Figure 6-A, iv-vi and B) and HA::IER3IP1^L78P^ (Figure 6-A, vii-iv and B) show a partial colocalization, whereas, HA::IER3IP1^T79*^ does not show any significant co-localization with Rab11 (Figure 6-A, x-xii and B). Next, we checked whether the mutant proteins interact with Rab11. We performed an immunoprecipitation assay using anti-HA on extracts from cells expressing HA::IER3IP1^Ref^, HA::IER3IP1^V21G^ and HA::IER3IP1^L78P^ but we find that interaction between mutant proteins HA::IER3IP1^V21G^ or HA::IER3IP1^L78P^ and Rab11 is reduced (Figure 6-C). These data confirm that disease-associated mutations affect IER3IP1 localization to Rab11 vesicles and interaction with Rab11. Overall, our data indicate that the loss of IER3IP1 affects Rab11 vesicle docking/tethering or fusion to the ingressing furrow during cytokinesis and suggest that IER3IP1 may be working along with Rab11 to regulate vesicle docking/tethering or fusion.

**Figure 6:**
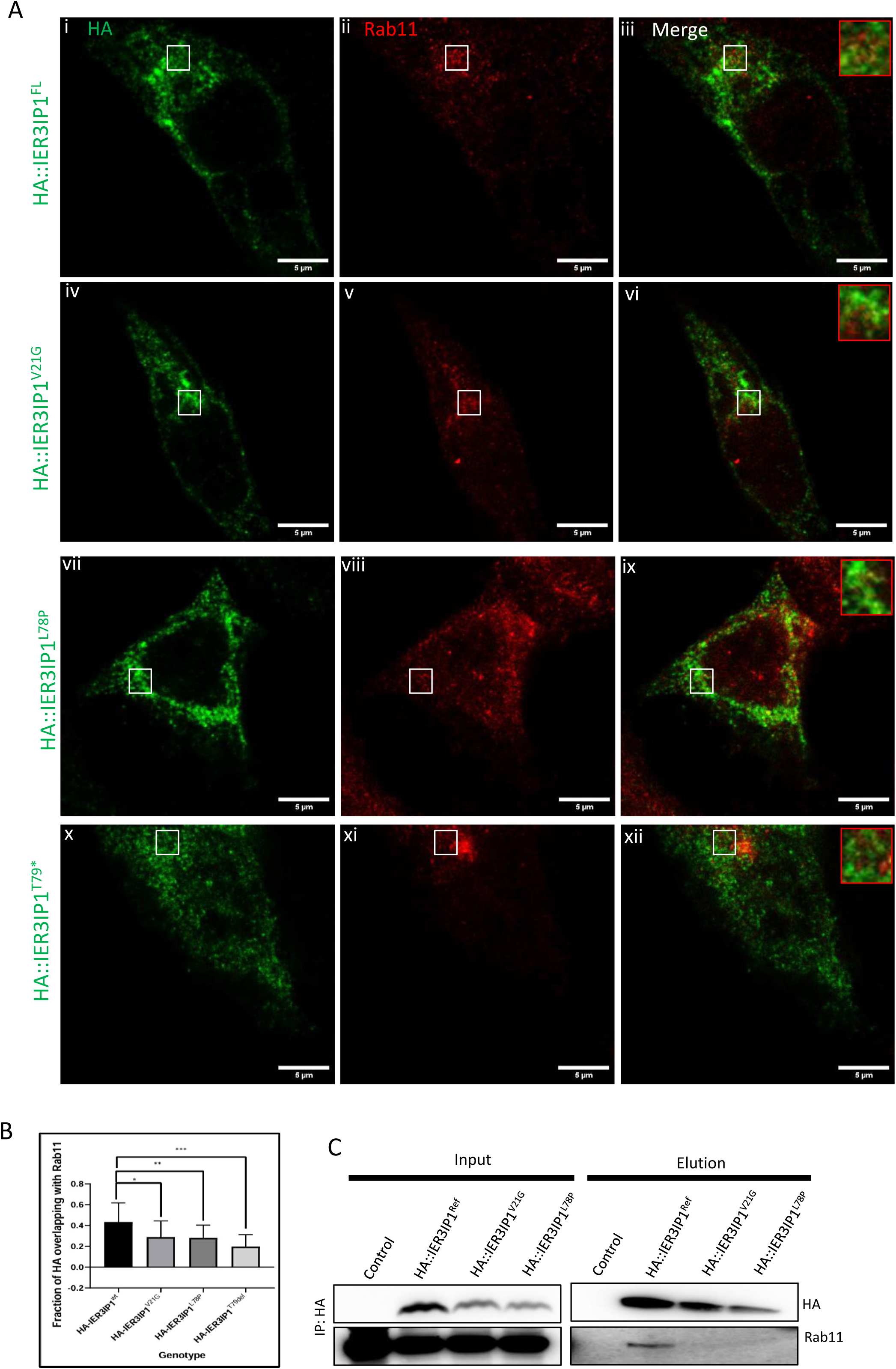
Disease associated mutations perturb IER3IP1 localization to Rab11 vesicles. A. HCT116 cells expressing *IER3IP1ref cDNA* (i-iii) and disease associated variants *V21G* (iv-vi) , *L78P* (vii-ix) and *T79\** (x-xii) were stained with anti HA and Rab11 antibodies, red insets shows the magnified view of the white inset in the same panel, scale bar=5µm. B. shows Mander’s correlation coefficient for fraction of HA overlapping with Rab11 for HA-IER3IP1^Ref^ ,HA-IER3IP1 ^V21G^,HA-IER3IP1 ^L78P^ and HA-IER3IP1 ^T79*^, ∗∗p<0.01, ∗∗∗p<0.001, ∗∗∗∗p<0.0001. C. Immunoprecipitation assay showing HA::IER3IP1^Ref^ , HA-IER3IP1^V21G^ and HA-IER3IP1^L78P^ were immunoprecipitated from HCT116 cells lysates using anti-HA beads and probed with anti-HA and anti-Rab11 antibodies.

## Discussion

In this study, we characterized *Inseparable*, the fly orthologue of *IER3IP1*, and shed light on its role in brain development and vesicle trafficking. We show that *Insep* is expressed in larval brains (Figure S1-B), and its loss during development leads to early larval lethality and small brain phenotype (Figure 1-B, C and D), a phenotype resembling microcephaly in humans. We found that *Insep* deficient NBs fail to complete cytokinesis, despite the normal actomyosin ring assembly and initial constriction. Failures in completing cytokinesis results in limiting neurogenesis that leads to microcephaly in flies. We show that human *IER3IP1* can rescue the phenotypes associated with *Insep* loss, indicating that the biological function of *Insep* is conserved in humans (Figure 1-D).

Similar to NBs, loss of *IER3IP1* in HCT116 and HEK293 cells also results in cytokinesis failure (Figure 4-A and Figure S5-B). *Insep* deficient NBs and *IER3IP1* deficient HCT116 cells that failed to complete cytokinesis accumulate Rab11 vesicles near the site of furrow ingression or cell equator (Figure 3 and Figure 4-B). Studies in several model systems have shown that Rab11 vesicles travel to the advancing furrow and fuse with them to complete cytokinesis and abscission (38–45). In *Insep* deficient NBs and *IER3IP1* depleted HCT116 cells, Rab11 vesicles travel to and accumulate near the advancing furrow and midbody (Figure S4 and Figure 4), suggesting that their transport to these locations is not affected; however, their fusion with the advancing furrow may be affected. This is in agreement with the previous reports where it was shown that defects in vesicle tethering and fusion cause an abnormal accumulation of Rab11 vesicles near the cleavage furrow in *Drosophila* (38, 46) *S. cerevisiae* (47) *S. pombe* (48) and at the midbody in human cells (49). Our data suggests that the assembly of the actomyosin ring and initial constriction is not affected in *Insep* deficient NBs; however, lack of Rab11 vesicles fusion to the ingressing furrow may cause destabilization of actomyosin ring at late stages of cytokinesis eventually leading to furrow regression and cytokinesis failure. Vesicle fusion to the ingressing furrow has been shown to be essential for the stability and constriction of the actomyosin ring during cytokinesis (38, 41, 50, 51). Beside cytokinesis failure, we also observed several vesicles accumulated beneath the plasma membrane in *Insep* deficient NBs (Figure 3-B) suggesting that vesicle tethering/ fusion to plasma membrane may also be affected.

We found that Insep and IER3IP1 localize to Rab11 vesicles and interact with Rab11 (Figure 5). Numerous studies have shown that Rab11 is involved in vesicle docking/ tethering and fusion to the plasma membrane (52–55). Rab11 interacts with vesicle docking factor Munc13-4 that helps in priming and docking of vesicles at the plasma membrane (53, 56). Rab11 has also been shown to interact with the exocyst complex that helps in tethering vesicles to plasma membrane (54, 57). In addition, loss of tethering, and fusion complex proteins results in an abnormal accumulation of vesicles (46, 54, 58). These reports and our observations that Insep and IER3IP1 localize to Rab11 vesicles, interact with Rab11, and their loss leads to an accumulation of Rab11 vesicles together strongly support that Insep/IER3IP1 is working along with Rab11 in vesicle docking/tethering or fusion and its role in these processes is indispensable.

To understand the pathogenic mechanism linked to human *IER3IP1* alleles, we analysed HA-IER3IP1^V21G^, HA-IER3IP1^L78P^, and HA-IER3IP1^T79*^ proteins for their stability and localization to ER and Rab11 vesicles. We found that these mutations decrease the stability of the IER3IP1 protein (Figure S1-C and Figure S5-E). A similar observation was reported for disease-linked A18V variant of *IER3IP1*(59). In our localization analysis, we found that the V21G variant does not affect the localization of IER3IP1 to ER. The L78P variant severely reduces the localization of IER3IP1 to ER, and the T79* variant does not show any significant localization to ER (Figure 5-D). The L78P mutation affects the C-terminal TM domain, which has been shown to be essential for IER3IP1 localization to ER (28, 29); this mutation may alter the TM domain structure resulting in reduced localization to ER. The T79* frameshift mutation leads to the expression of a truncated protein that does not have the C-terminal TM domain, therefore no localization to ER was observed. We also find that V21G, T79* and L78P variants affect the localization of IER3IP1 to the Rab11 vesicles, the V21G variant shows partial localization, whereas, the T79* and L78P variants do not show any significant localization to the Rab11 vesicles. We also find that the V21G and L78P variants reduce IER3IP1 binding to Rab11 (Figure 6-A and B). Our observations along with *Insep* rescue data (Figure 1-D) suggest that V21G, T79* and L78P variants are severe loss of function mutations.

In yeast, Yos1P localizes to COPII vesicles and its loss leads to ER to Golgi transport defects. A recent report showed that IER3IP1 interacts with TMEM167A, a Golgi membrane-localized protein, and suggested that it may be facilitating the docking or fusion of COPII vesicles with the Golgi membrane (59). Our data show that IER3IP1 localizes to Rab11 vesicles, interacts with Rab11, and may be involved in Rab11 vesicle fusion to the advancing cleavage furrow during cytokinesis. In addition, human disease-associated mutations abolish IER3IP1 binding to TMEM167A (59) and Rab11 (our data). These observations suggest that IER3IP1 may play a role in vesicle docking/tethering or fusion at various stages of the secretory pathway. However, the precise molecular function of *IER3IP1* during vesicle docking/tethering or fusion remains to be determined.

## Materials and Methods

### *Drosophila* stocks and husbandry

Fly stocks used in this study are listed in Table 1. All flies were maintained at 25°C and grown on standard cornmeal and molasses medium. Crosses were performed at temperature indicated (25°C or 29°C).

**Table 1.** Fly stocks used in this study.

| **Fly strains** | **Source** |
| --- | --- |
| w<sup>1118</sup>; P{w<sup>+</sup>mC}=UAS-EGFP}5a.2 | BDSC |
| y<sup>1</sup> M{w<sup>+</sup>mC}=nanos-Cas9.P}ZH-2A w<sup>*</sup> | BDSC |
| y<sup>1</sup> M{w<sup>+</sup>mC}=Act5C-Cas9.P}ZH-2A w<sup>*</sup> | BDSC |
| y<sup>1</sup> sc<sup>*</sup> v<sup>1</sup> sev21; P{TKO.GS01105}attP40 | BDSC |
| y<sup>1</sup> sc<sup>*</sup> v<sup>1</sup> sev21; P{TRiP.HMS05814}attP40 | BDSC |
| w<sup>1118</sup>; P{GD3868}v9202 | VDRC |
| w<sup>*</sup>; P{w<sup>+</sup>mW.hs}=GawB}insc[Mz1407] | BDSC |
| y<sup>1</sup> w<sup>*</sup>; P{w<sup>+</sup>mC}=Act5C-GAL4}/TM6B, Tb<sup>1</sup> | BDSC |
| w<sup>*</sup>; P{w<sup>+</sup>mC}=sqh-mCherry.M}3 | BDSC |
| y<sup>*</sup>w<sup>*</sup>; Inseperable3XP3-GFP/ TM3, Sb,Ser | This study |
| y<sup>*</sup>w<sup>*</sup>; InseperableT2A>GAL4/ TM3, Sb,Ser | This study |
| y<sup>*</sup>w<sup>*</sup>; P{UAS-Inseperable}3 | This study |
| y<sup>*</sup>w<sup>*</sup>; P{UAS-HA-Inseperable}3 | This study |
| y<sup>*</sup>w<sup>*</sup>; P{UAS-Inseperable-HA}3 | This study |
| y<sup>*</sup>w<sup>*</sup>; P{UAS-IER3IP1}3 | This study |
| y<sup>*</sup>w<sup>*</sup>; P{UAS-HA-IER3IP1Wt}3 | This study |
| y<sup>*</sup>w<sup>*</sup>; P{UAS-HA-IER3IP1V21G}3 | This study |
| y<sup>*</sup>w<sup>*</sup>; P{UAS-HA-IER3IP1L78P}3 | This study |

### Plasmid construction

Oligonucleotides and plasmid used are detailed in Table 2 and Table 3, respectively.

**Table 2.**
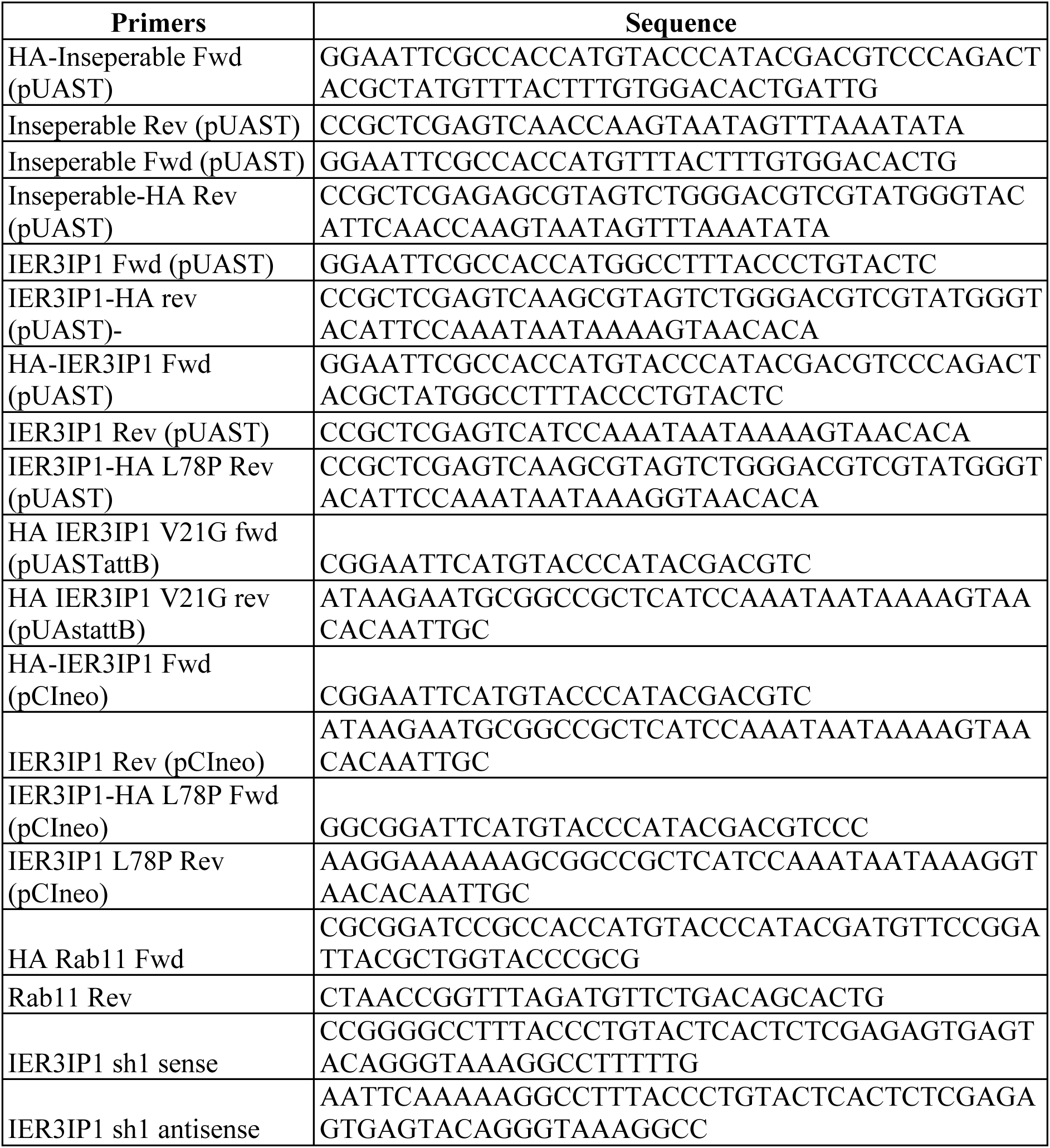
Oligoes/Primers used in this study.

**Table 3.** Plasmids used in this study.

| Plasmid | Source |
| --- | --- |
| pUASTattB | DGRC, 1419 |
| pUASTattB-Inseperable | This study |
| pUASTattB -HA-Inseperable | This study |
| pUASTattB -Inseperable-HA | This study |
| pUASTattB -IER3IP1 | This study |
| pUASTattB -HA-IER3IP1 | This study |
| pUASTattB -IER3IP1-HA | This study |
| pUASTattB -IER3IP1-HA V21G | This study |
| pUASTattB -IER3IP1-HA L78P | This study |
| pCIneo | Addgene, E1841 |
| pCFD3-dU63gRNA | Addgene, 49410 |
| pLentiDest | Addgene, 13320 |
| pLKO.1 TRC | Addgene, 10878 |
| mCherry-Rab11a | Addgene, 55124 |
| pAdTrack-CMV | Addgene, 16405 |

pUC57-Insep^3XP3-GFP^. Construct pUC57-Insep^3XP3-GFP^ was commercially synthesized (Genscript). It contains 1Kb Left homology arm of IER3IP1 followed by a gene trap cassette attB-3XP3-EGFP-SV40pA-attB and 1 KB Right homology arm of IER3IP1.

pCFD3-dU63-Insep-gRNA#1 and pCFD3-dU63-Insep-gRNA#2: The sense and antisense oligos for the guide RNA were phosphorylated (37°C for 30 mins), heated at 95°C and annealed by slow cooling at the rate of 0.1°C/sec to 25°C in a thermocycler. Annealed oligos were ligated into the pCFD3 vector using Bbs1 sites.

pUASTattB-HA-Inseparable and pUASTattB-Inseparable-HA. Full length Inseparable cDNA fragment was commercially synthesized (Genscript). Fragments: HA-Inseparable and Inseparable-HA were amplified using the primers listed in Table 2, digested and cloned into the pUASTattB vector between EcoRI and XhoI sites.

pUASTattB-IER3IP1, pUASTattB-HA-IER3IP1, pUASTattB-HA-IER3IP1^v21G^ and pUASTattB-HA-IER3IP1^L78P^. A plasmid containing full length IER3IP1 (clone ID-TOLH-1517029) was obtained from Transomic technologies. Inserts full length IER3IP1, HA-IER3IP1 and HA-IER3IP1^L78P^ were amplified using primers listed in Table 2, digested and cloned into the pUASTattB vector between EcoRI and XhoI or NotI. The fragment HA-IER3IP1^V21G^ was commercially synthesized (Genscript) and cloned into pUASTattB between EcoRI and XhoI or NotI.

pCIneo-HA-IER3IP1, and pCIneo-HA-IER3IP1^L78P^. HA-IER3IP1 was digested from pUAST constructs and subcloned into pCIneo between EcoRI and NotI. Fragment HA-IER3IP1^L78P^ was amplified using primers listed in Table 2 and cloned into the pCIneo between EcoRI and NotI.

pCIneo-HA-IER3IP1^V21G^ and pCIneo-HA-IER3IP1^T79*^. The fragments HA-IER3IP1^V21G^ and HA-IER3IP1^T79^ were commercially synthesized (Genscript) and cloned into pCIneo between EcoRI and NotI.

plko.1trc-IER3IP1-shRNA#1. Forward and reverse Oligos were designed against the shRNA target sequence (5’ GGCCTTTACCCTGT 3’) with EcoR1 and Age1 overhangs, respectively. Oligos were heated at 70°C for 4 mins followed by slow cooling at the rate of 0.1°C / sec to 25°C. Annealed oligos were ligated into the plko.1trc vector between EcoR1 and Age1 sites.

### Generation of *Insep* T2A-GAL4, null mutant and transgenic lines

***Insep^T2A-GAL4^*:**Construct pBSK-Insep^T2A-GAL4^ was co-injected with pCFD3-dU63-Insep-gRNA#1 and pCFD3-dU63-Insep-gRNA#2 into *(y[1] M{w[+mC]=nanos-Cas9.P}ZH-2A w[*]* embryos (54591, BDSC).

Injected adults were crossed to w[1118]; P{w[+mC]=UAS-EGFP}5a.2 and progeny were screened for EGFP expression and individual *Insep^T2A-GAL4^* lines were established.

***Insep^3XP3-EGFP^*:** Construct pBSK-Insep^3XP3-GFP^ was co-injected with pCFD3-dU63-Insep-gRNA#1 and pCFD3-dU63-Insep-gRNA#2 into *(y[1] M{w[+mC]=nanos-Cas9.P}ZH-2A w[*]* embryos (54591, BDSC). Injected adults were screened for the expression of EGFP in their eyes. Deletion mutants were selected based on the expression of EGFP in their eyes.

***UAS-Insep* and *UAS-IER3IP1* transgenic lines:** Constructs pUAST-HA-Inseparable, pUAST-Inseparable-HA, pUASTattB-IER3IP1, pUASTattB-HA-IER3IP1, pUASTattB-HA-IER3IP1^v21G^ and pUASTattB-HA-IER3IP1^L78P^ were injected in *y1w∗* ; *VK37* embryos. Injected adults were screened for transgenes with red eyes.

### Cell culture and transfections

Cell lines used in this study are listed in Table 4.

**Table 4.** Cell lines used in this study.

| Cell line | Source |
| --- | --- |
| HCT116 cells | ATCC, ATCC-CCL-247 |
| HEK293 cells | ATCC, ATCC-CRL-1573 |
| 293T cells | ATCC-CRL-3216 |
| D. melanogaster: Cell line S2: S2-DRSC | DGRC, 181 |

Cell culture: HEK293 cells were cultured in complete DMEM (4mM L glutamine, 4500 mg/L glucose, 1mM sodium pyruvate, 1500 mg/L Sodium bicarbonate) supplemented with 10% fetal bovine serum, 100 U/ml penicillin, and 0.10 mg/ml streptomycin at 37°C with 5% CO2. HCT116 cells were cultured in McCoy’s 5A Medium w/ L-Glutamine and Sodium bicarbonate (Himedia, Cat# AL057A) supplemented with 10% fetal bovine serum, 10% MEM Non Essential Amino Acids Solution (Himedia, Cat # ACL006), 100 U/ml penicillin, and 0.10 mg/ml streptomycin at 37°C with 5% CO_2_. *Drosophila* S2 cells were cultured in Schneider’s Insect Medium (Sigma) supplemented with 0.4 g/L sodium bicarbonate, 0.6 g/L calcium chloride,10 % heat-inactivated FBS, 100 U/ml penicillin, and 0.10 mg/ml streptomycin at 25°C with no requirement of CO_2._

Transient transfection: HCT116/ HEK293 cells were cultured in a 6-well plate or 10 cm petri dish or 15 cm petri dish. At 60% confluency the cells were transiently transfected with plasmid DNA (2.5-3.5 µg for a 6-well plate, 20 µg for a 10 cm petri dish and 50 µg for a 15 cm petri dish) using Lipofectamine 3000 (Invitrogen, L3000015) or polyethylenimine (Polysciences, Cat # 24765). S2 cells were transfected with *UAS-Inseparable-HA* and *Act GAL4* plasmids at 60-70% confluency. Transfections were performed using Effectene transfection reagent (Qiagen,Cat # 301425). After transfection, the cells were grown for 24 hours and then processed for immunoprecipitation.

### Lentiviral particles preparation and IER3IP1 knock-down in cells

Lentiviral particle production: HEK293T (ATCC-CRL-3216) cells were cultured in complete DMEM (4mM L glutamine, 4500 mg/L glucose, 1mM sodium pyruvate, 1500 mg/L Sodium bicarbonate) supplemented with 10% fetal bovine serum, 100 U/ml penicillin, and 0.10 mg/ml streptomycin in 5% CO2 at 37°C. The pLentiDest-IER3IP1 shRNA#1 construct along with the three packaging vectors p-VSVG (Addgene 8454), pRSV (Addgene 12253), pMDL (Addgene 12251)) were transiently transfected into HEK293T cells using Lipofectamine 3000 and OPTI MEM media (Thermo Fischer Scientific, 31985070). Media was changed after 15 hours. Thereafter, the lentivirus containing supernatant was harvested and concentrated using Lenti X concentrator (Takara, 631231) after 24 hours. For the control viral particles production, pLentiDest containing a scrambled sequence was used.

IER3IP1 knockdown: HCT116 and HEK293 cells were seeded at 20% confluency and transduced with the IER3IP1 shRNA viral particles and Scramble shRNA (control) using hexadimethrine bromide (8µg/ml) to improve efficiency of viral transduction. 24 hours later, transduced cells were selected using Puromycin (6 µg/ml for HEK293 and 35 µg/ml for HCT116) over a period of two days. A non-transduced control treated with puromycin was used to ensure accurate selection of transduced cells. Post selection, the knockdown was confirmed using western blotting.

### Nocodazole treatment

Cells were treated with 100 ng/µl Nocadozole (Sigma, Cat # 31430-18-9) for 16 hours. Post treatment, the cells were rinsed three times with 1X PBS and released with puromycin containing media for 2 hours to capture anaphase and early telophase and 15 hours to enrich cells with cytokinesis defect.

### Immunocytochemistry

Primary and secondary antibodies are listed in Table 5

**Table 5.**
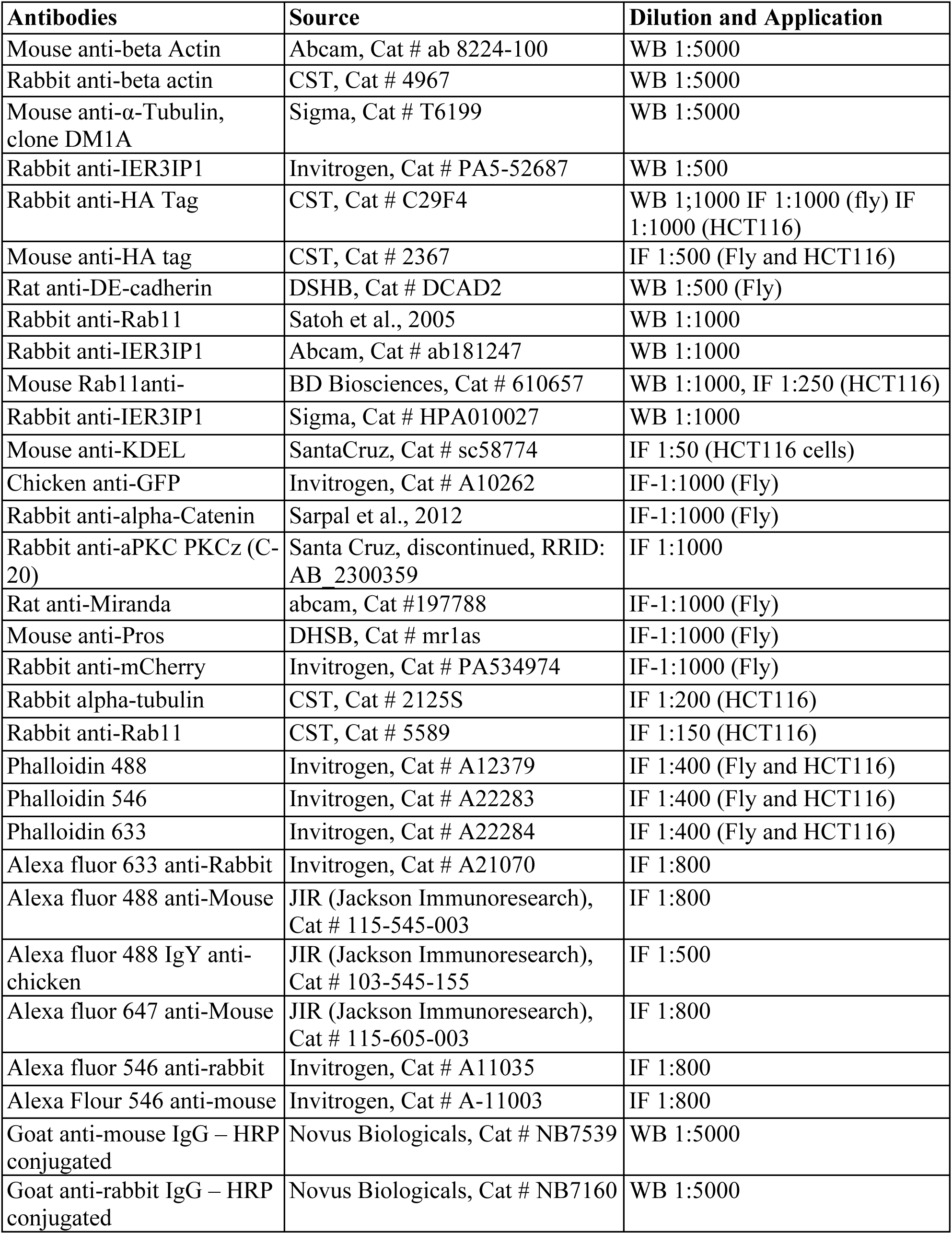
Antibodies used in this study.

Larval brains were dissected in PBS, fixed with 3.7% PFA for 20 minutes at room temperature and washed with PBST (PBS containing 0.3% Triton-X). Fixed brains were blocked in PBST containing 10% normal goat serum for 1 hour at room temperature (RT) and incubated in primary antibodies diluted in blocking solution overnight at 4°C. Samples were washed with PBST, incubated with the secondary antibodies for 2 hours at RT and mounted in 70% glycerol or ProLong™ Gold Antifade mounting media (Invitrogen, P36934). Imaging was done using Zeiss LSM 880 and Leica TCS SP8 confocal microscopes. STED imaging was done using a Leica TCS STED microscope.

HCT116 and HEK293 cells were seeded on coverslips in a 6-well plate and post shRNA or Nocodazole treatment were washed with PBS, fixed with 3.7% Paraformaldehyde for 15 minutes at RT. Further, the cells were permeabilized with PBST (PBS/ 0.2% Triton-X) at RT for 15 minutes. Cells were blocked in 2% BSA in PBST for 1 hour at RT and incubated with primary antibodies diluted in the blocking solution overnight at 4°C. Cells were washed 3 times with PBST at RT for 5 minutes and incubated with secondary antibodies diluted in the block solution for 2 hours at RT. The cells were washed with PBST for 5 minutes at RT and stained with DAPI (1 μg/ml) (without dapi for STED samples) for 5 minutes at RT followed by a final wash of PBST for 10 minutes at RT. Cells were then mounted using Prolong Gold antifade mounting media. The slides were imaged by confocal microscopy on Zeiss LSM 880 and Leica TCS SP8 confocal microscope. STED imaging was done using Leica TCS STED microscope

### Immunoblotting and co-immunoprecipitation

Immunoblotting: HCT116/HEK293 cells were harvested and lysed in a lysis buffer (50mM Tris,150mM NaCl, 0.2% Triton X 100) with Protease and Phosphatase Inhibitors by incubating on ice for 30 minutes. After lysis, the samples were centrifuged at 4°C for 10 mins at max speed and supernatants were transferred to new tubes. Total proteins were estimated using the BCA kit (Thermo Scientific, 23225), separated on SDS-PAGE and blotted on PVDF or Nitrocellulose membrane. The membrane was blocked in blocking buffer either 5% BSA (Himedia Cat # MB083) or 5% Blotto (Santacruz Cat # sc-2325 for) for 1 hr at RT. Blots were incubated with primary antibodies at appropriate concentrations overnight at 4°C, washed with TBST (TBS/0.1% Tween20) three times for 10 minutes each and incubated with secondary antibodies for 2 hours at RT. Blots were developed using SuperSignal West Pico PLUS Chemiluminescent substrate (Thermo Scientific, Cat.No. 34579) on a Vilber Lourmat Chemidoc and Biorad Chemidoc.

Immunoprecipitation: HCT116 cells were seeded in 10 cm or 15 cm petri dishes. At 60% confluency, cells were transfected with constructs expressing HA::IER3IP1^Ref^ and mutant variants using Lipofectamine 3000 or polyethyleneimine (Polysciences, 24765) and cultured for 48 hours. Cells were pelleted and lysates were prepared in 100 µl Lysis Buffer (50 mM Tris pH8.0, 100 mM NaCl, 5mm EDTA, 1% Digitonin, 1x protease and phosphatase inhibitors) , incubated on ice for 30 mins and centrifuged for 15 min at 4°C. Lysates were diluted with 800µl wash buffer (50 mM Tris pH8.0, 100 mM NaCl, 5mm EDTA) and incubated with HA beads (EZview™ Red Anti-HA Affinity Gel, Merck, E6779) or RFP beads (ChromoTek, rta-20) overnight at 4°C on rocker. The beads were washed with the wash buffer for three times 5 minutes each. HA tagged proteins were eluted with HA peptide (Sigma, 12149 -final concentration 100ug/ml, 4 hours’ incubation at 4°C or with HA peptide (Thermo, 26184, final concentration 0.1mg/ml, 2 hours’ incubation at 37°C) or by boiling the beads at 95°C with 50 ul of 2x Laemmli buffer. The mCherry tagged proteins were eluted by boiling the beads at 95°C with 50ul of 2X Laemmli Buffer for 10 minutes. Untransfected HCT116 cells were used as control.

The transfected S2 cells were washed with 1X PBS and centrifuged at 1500 rpm for 3 minutes. The lysates were prepared in 100 µl Lysis Buffer (50 mM Tris pH 8.0, 100 mM NaCl, 5mm EDTA, 1% Digitonin, 1x protease and phosphatase inhibitors), incubated on ice for 30 mins and centrifuged for 15 min at 4°C. Lysates were diluted with 800µl wash buffer (50 mM Tris pH8.0, 100 mM NaCl, 5mm EDTA) and incubated with HA beads (EZview™ Red Anti-HA Affinity Gel, Merck, E6779) overnight at 4°C on rocker. The beads were washed with the wash buffer for three times 5 minutes each. HA tagged proteins were eluted with HA peptide (Sigma, 12149 -final concentration 100ug/ml, 4 hours’ incubation at 4°C or with HA peptide (Thermo, 26184, final concentration 0.1mg/ml, 2 hours’ incubation at 37°C) or by boiling the beads at 95°C with 50 ul of 2x Laemmli buffer. Untransfected S2 cells were used as control.

### TEM

For TEM, larval brains were dissected in an ice cold PB buffer pH7.4 and fixed in 2% glutaraldehyde (made in 0.1M cacodylate buffer pH 7.2) for 2 hours at RT and later at 4°C overnight. Samples were washed with 0.1M cacodylate buffer for 5 min and incubated with 1% osmium tetroxide for 2 hours at RT in dark conditions. Thereafter, they were once again washed with 0.1M cacodylate buffer for 5 min and dehydrated using increasing concentrations of ethanol – 20%-50%-70% (twice for 10 min each). Samples were then stained with 1% uranyl acetate in 70% ethanol for 2 hours at 4°C in dark conditions and stored at 4°C overnight. The next day, samples were dehydrated in 95% and 100% ethanol (thrice for 5 min each). Samples were then treated with 100% polypropylene oxide (three times for 5 min each) followed by Resin infiltration gradually using epoxy resin prepared in polypropylene oxide ratio 1:1 and 2:1 for 1 hour each at RT. Thereafter, pure resin infiltration (100% epoxy) was done for 3 hours at RT. Samples were then transferred to fresh resin (100% epoxy) and allowed to polymerize at 60°C for 48 – 72 hours. Samples were sectioned as 70nm thin sections and stained as follows: 2% uranyl acetate for 30 mins in dark at RT, washed in milli-Q water and then stained with lead citrate for 8 mins followed by washing in milli-Q water and dried overnight at RT. Imaging was done using Talos 120C Transmission Electron Microscope and images captured at 120kV using Ceta camera in Velox software.

### Quantification and Statistical Analysis

#### Brain Volume

68 hrs AEL brains from *Insep CN* and 45 hrs AEL brains from *Insep^N^* were stained and imaged using LSM 800 confocal system. One lobe from each brain was imaged and a total of 10-15 brains were analyzed per genotype. Resulting stacks were analyzed using ImageJ software to compute the brain volume. At first, the area of each stack was measured. It was multiplied with the thickness to obtain the volume of one stack. Next the volume of all the stacks was added to obtain the volume of the entire lobe. This method was followed for all the brains. The data is plotted by taking the average of total brain lobe volume (μm ^3^) for all brains. Statistical significance was determined using t-test.

#### Cytokinesis defect

68 hrs AEL brains from *Insep CN* and 45 hrs AEL brains from *Insep^N^* were stained and imaged using LSM 800 confocal system. Total number of NBs and total number of multinucleated NBs were manually counted from the stacks using ImageJ. The data is plotted as a percentage of multinucleated NBs present per genotype.

#### Co-localization analysis

We used Mander’s correlation coefficient to measure co-localization between HA:Insep and Rab11 for third instar larval brains and HA:IER3IP1-Rab11 for HCT 116 cells. The fraction of HA:Insep/HA:IER3IP1 signal overlapping with Rab11 and Rab11 signal overlapping with HA:INSEP/HA:IER3IP1 has been plotted in the graphs. The ImageJ software was used for this analysis by installing the JACoP plugin. To compare the co-localization between HA::IER3IP1^Ref^ HA-IER3IP1^V21G^/HA-IER3IP1^L78P^/HA-IER3IP1^T79*^and Rab11, similarly between HA::IER3IP1^Wt^ HA-IER3IP1^V21G^/HA-IER3IP1^L78P^/HA-IER3IP1^T79*^ and KDEL; the fraction of HA signal overlapping with Rab11/KDEL was computed using the JACoP plugin .Statistical significance was determined using t-test.

## Acknowledgements

We thank Hugo J Bellen, Manish Jaiswal and Richa Ricky for various fly lines used in this study. We are grateful to Donald F Ready and Hongyan Wang for sharing antibodies. Stocks from the Bloomington *Drosophila* Stock Center and Vienna *Drosophila* Stock Center were used for this study. We thank *Drosophila* injection facility at Bangalore LifeScience Cluster, NCBS-TIFR for embryo injections. We are grateful to Deepa Balasubramanian and Manish Jaiswal for helping us in transgenic fly lines generation. We thank Mayukh Banerjee, Krittika Biswas, Deyasini Chakraborty and Andrea Johnson for technical help. We thank Janesh Kumar for troubleshooting the co-IP experiment. We thank the fly facility, microscopy facility and cell culture facility at CSIR-CCMB. SN-J started the gene expression screen in the lab of Hugo J Bellen-we thank his generous support. We thank Oguz Kanca, Santosh Kumar, Varun Choudhury and Manish Jaiswal for critical reading and editing of the manuscript and SN-J lab for fruitful discussion. AK is supported by DBT-SRF, SG is supported by CSIR-SRF, RV is supported by DBT/Wellcome India alliance (IA/I/18/1/503629). SN-J is supported by DBT/Wellcome India alliance (IA/I/18/1/503629), CRG-SERB (CRG/2022/007401) and CSIR-CCMB funds.

## Author Contributions

A.A.K. and S.N-J. designed the study; A.A.K., S.G., R.V., H.A., D.B. and S.N-J. performed the experiments; A.A.K., S.G., R.V., H.A., M.J. and S.N-J analyzed the data; and A.A.K, S.G., R.V., M.J. and S.N-J. wrote the manuscript.

---

**Figure S1.**
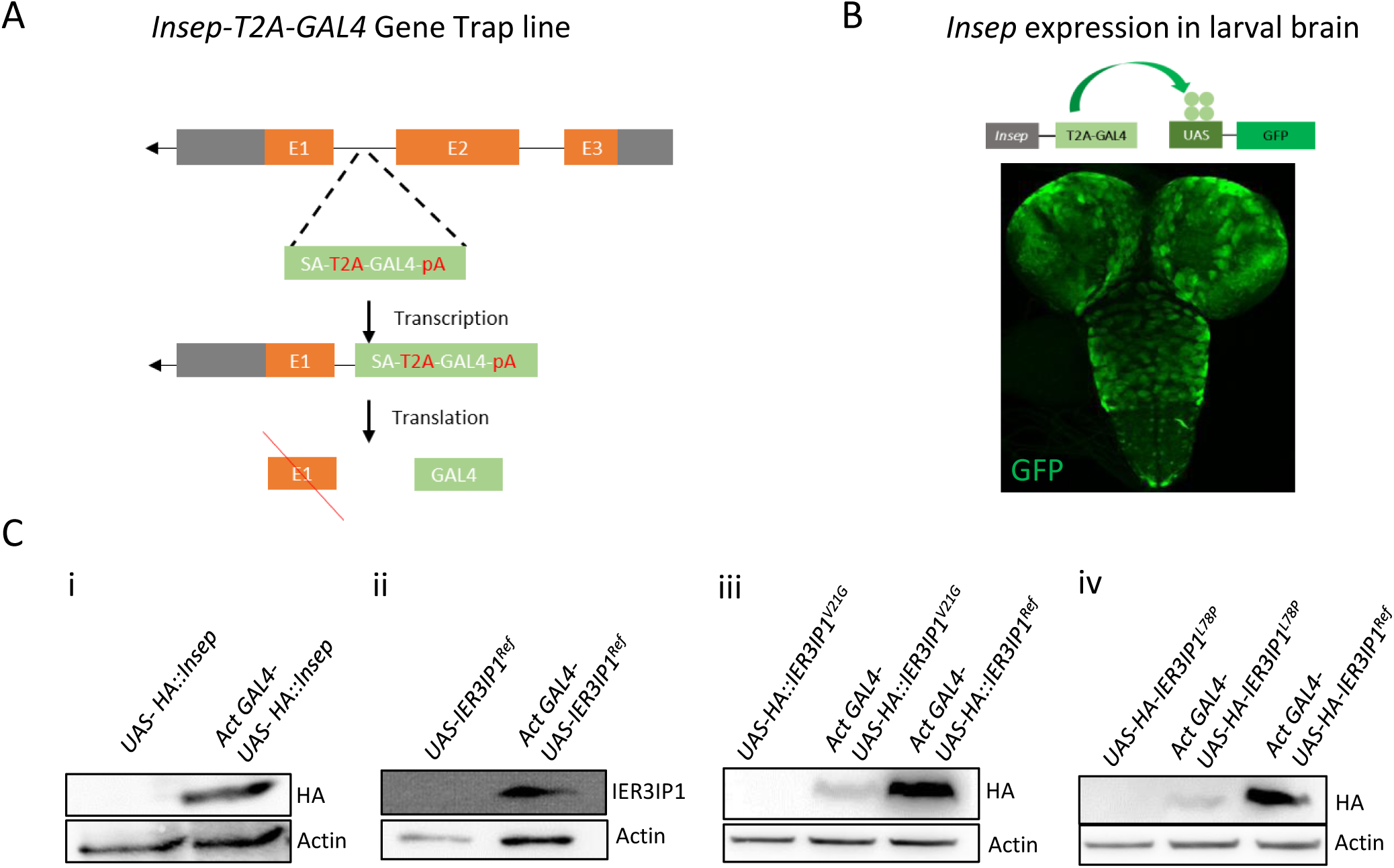
Expression of *Insep* and *IER3IP1* reference and disease associate variants in flies: A. Schematic diagram showing *Insep* gene locus, where the SA-T2A-GAL4-pA cassette is inserted in the first coding intron of *Insep* gene. The SV40 polyadenylation signal (pA) present in the cassette will cause a premature termination of transcription. During splicing, due of the Splice Acceptor (SA), the SA-T2A-GAL4 cassette will be in inserted into the mature RNA. Upon translation, the T2A sequence will cause truncation of INSEP protein and production of GAL4. B. *Insep-T2A-GAL4* gene trap driver line driving the expression of EGFP in L3 larval brain. C. Western blots of larval extract from (i) Control larvae *UAS-HA::Insep* (left) and larvae expressing *HA::INSEP* (right), (ii) Control larvae *UAS-IER3IP1^ref^* (left) and larvae expressing untagged *IER3IP1^Ref^* (right), (iii) Control larvae *UAS-IER3IP1^V21G^* (left), larvae expressing *HA::IER3IP1^V21G^* (middle) and *HA::IER3IP1^Ref^* (right), (iv) Control larvae *UAS-IER3IP1^L78P^* (left), larvae expressing *HA::IER3IP1^L78P^* (middle) and *HA::IER3IP1^Ref^* . Blots were probed with anti-HA (i, iii and iv), anti-IER3IP1 (ii) and anti-actin (loading control).

**Figure S2.**
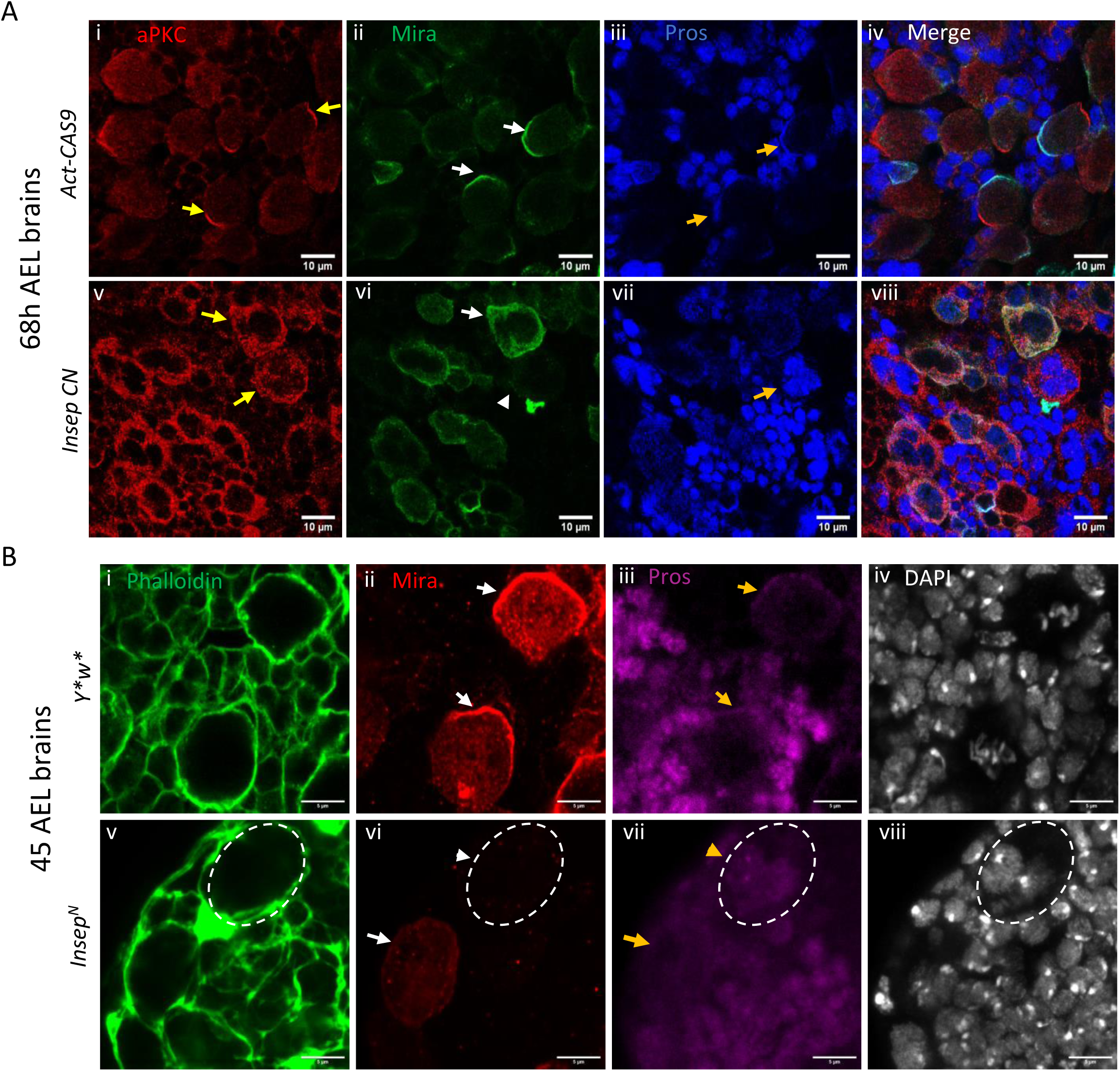
A. Confocal images of brain from *Act-Cas9* controls (n=9) and *Insep CN* (n=10) 68 AEL larvae stained with aPKC (red, i, iv, v and viii), Miranda (green, ii, iv, vi and viii), Pros (blue, iii, iv, vii and viii). Yellow arrows indicate localization of aPKC (i, iv, v and viii), white arrows indicate localization of Miranda (ii, iv, vi and viii), white arrow head indicates cell that has lost Miranda (vi), orange arrows indicate localization of Pros (iii and vii), scale bar=10µm B. Confocal images of brain from *y*w\** controls (n=11) and *Insep^N^* (n=10) 45 AEL larvae stained with Phalloidin (green, i and v), Miranda (red, ii and vi), Pros (magenta, iii and vii) and DAPI (gray, iv and viii). White arrows indicate localization of Mira (ii and vi), orange arrows indicate localization of Pros (iii and vii), white arrow head indicates cell that has lost Miranda (vi), orange arrowhead indicate localization of Pros (vii), scale bar=5µm. n = number of brains

**Figure S3:**
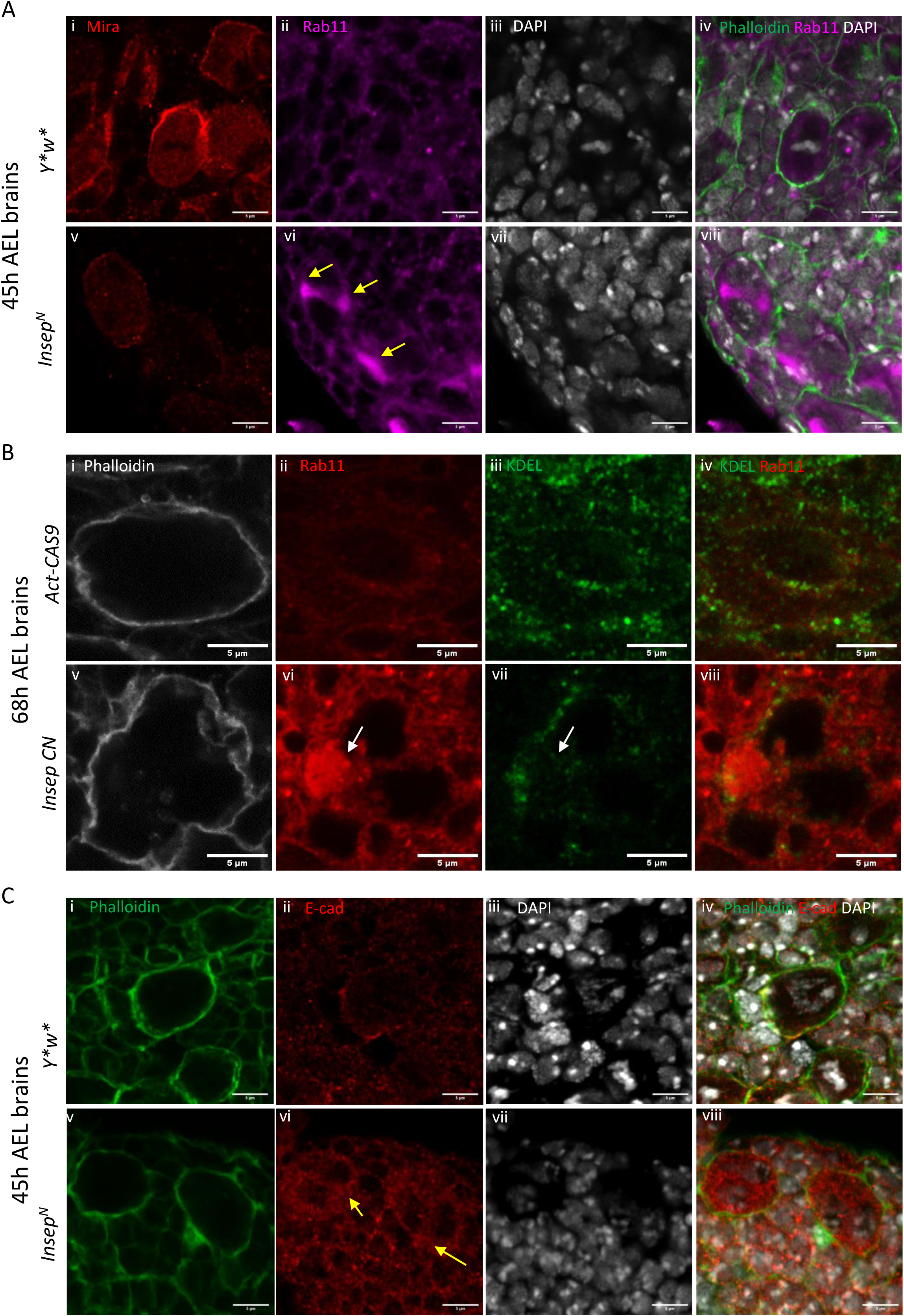
*Insep* mutant NBs accumulate Rab11 vesicle and it’s cargo DE-cad A. Confocal images of brain from 45h AEL *y*w\** controls (n=5) and *Insep^N^* (n=7) larval brains stained with Mira (red, i, and v), Rab11 (magenta, ii, iv, vi and viii), DAPI (gray, iii, iv, vii and viii), yellow arrows show accumulation of rab11 (vi). Scale bar=5µm B. Confocal images of brains from 68h AEL *Act-Cas9* controls (i-iv, n=11) and *Insep CN* (v-viii, n=17) stained with phalloidin (gray, i and v), KDEL (green, iii, iv, vii and viii) and Rab11 (red, ii, iv, vi and viii), white arrows shows Rab11 blob in (vii) and location in (vi). Scale bar=5µm. C. Confocal images of brain from *y*w\** controls (i-iv, n=11) and *Insep^N^* (v-viii, n=11) 45h AEL larvae stained with Phalloidin (green, i, iv, v and viii), DE-Cad (red, ii, iv, vi and viii), DAPI (gray, iii, iv, vii and viii), yellow arrows show accumulation of DE-Cad in *Insep CN* NB cytoplasm (vi). Scale bar=5µm

**Figure S4:**
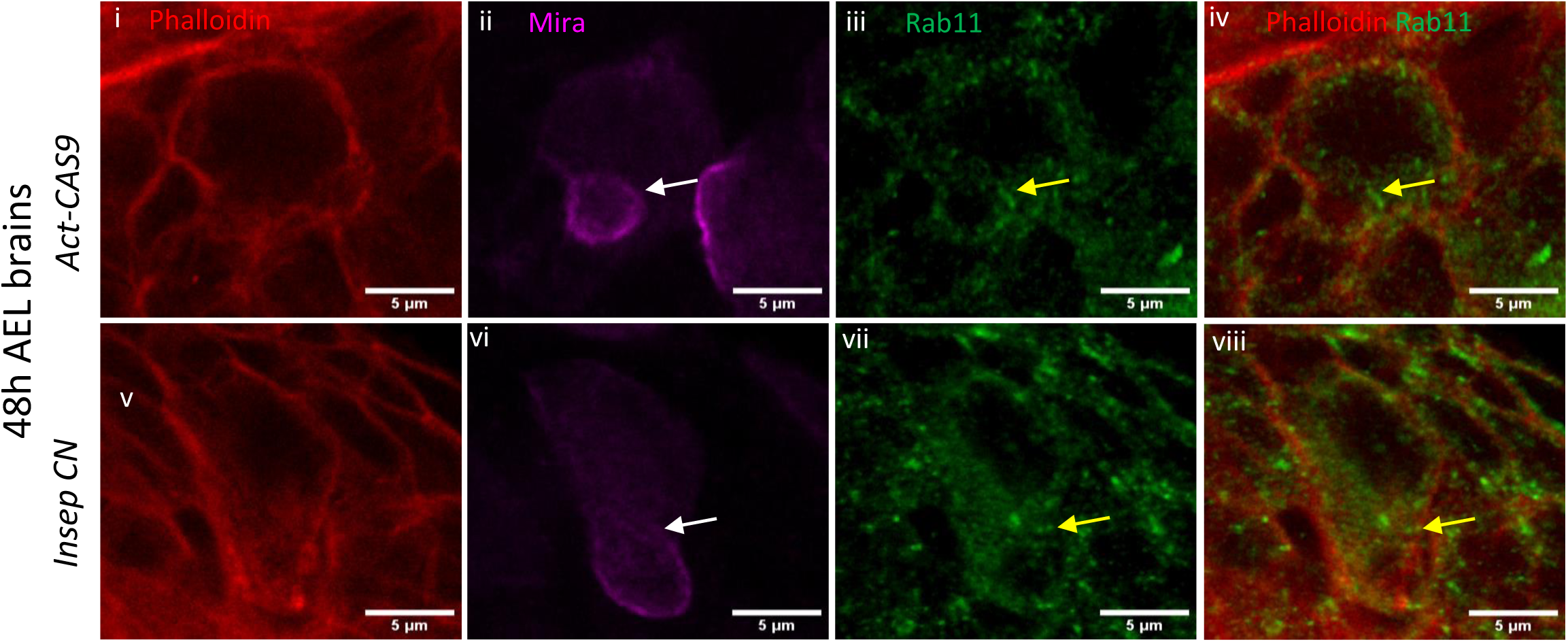
Confocal images of brain from *Act-Cas9* controls (i-iv, n=10) and *Insep CN* (v-viii, n=9) 48h AEL larvae stained with Phalloidin (red, i, iv, v and viii), Mira (magenta, ii, iv, vi and viii), Rab11 (green iii, iv, vii and viii), white arrows in (ii and vi) indicate the advancing furrow, yellow arrows in (iii and vii) show Rab11 vesicles assembled at the ingressing furrow. Scale bar=5µm

**Figure S5:**
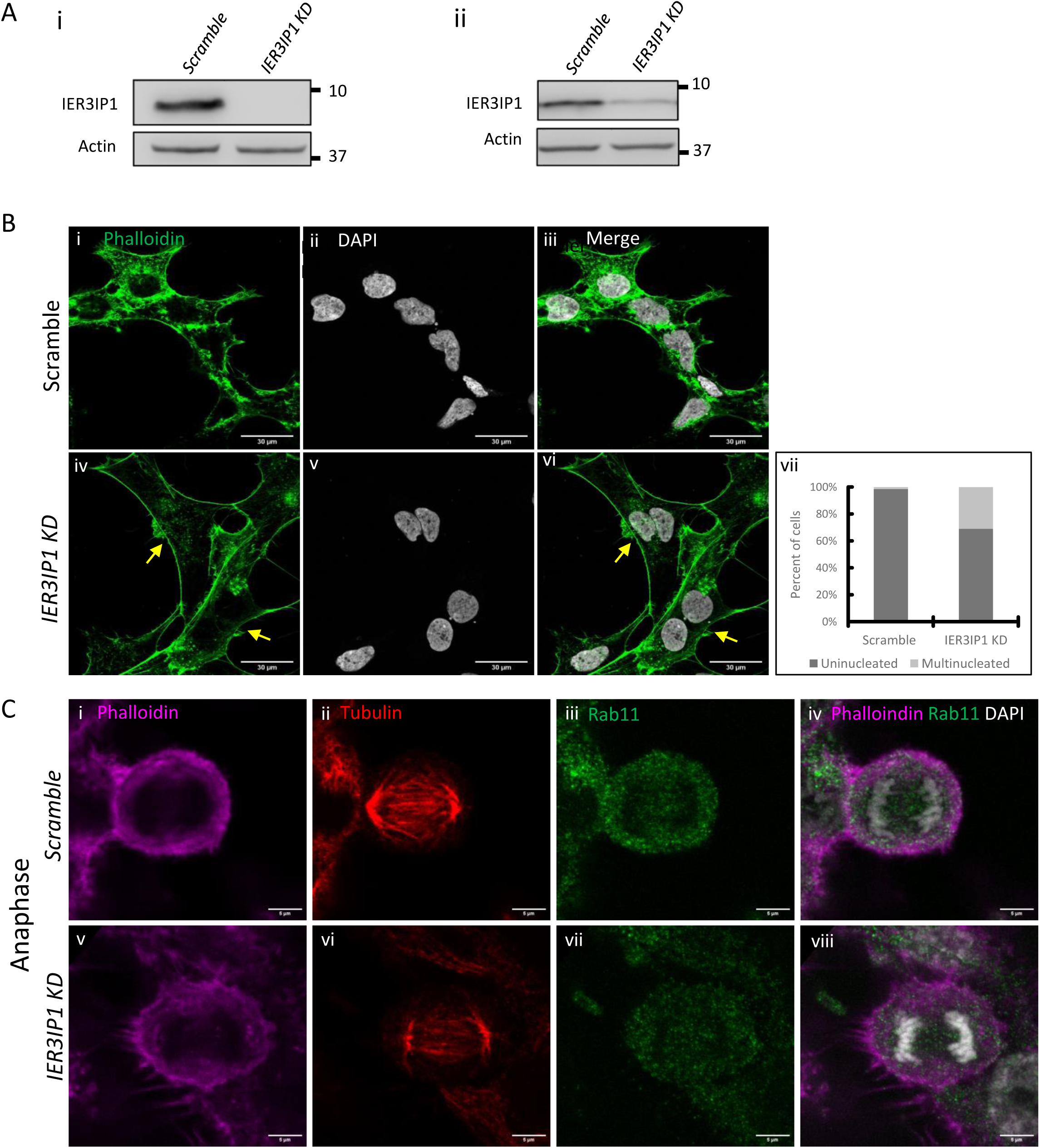
Loss of IER3IP1 leads to cytokinesis failure. A. Western blots showing shRNA mediated knockdown of *IER3IP1* in HCT116 (i) and HEK293 cells (ii), blots were probed with anti IER3IP1 antibody and actin (loading control) B. Confocal images of scramble control (i-iii) and *IER3IP1* shRNA treated HEK293 cells (iv-vi) stained with Phalloidin and DAPI. Yellow arrows in ( iv) and (vi) indicate cells with cytokinesis, vii shows quantification of (i-vi). Scale bar=10µm. C. Confocal images of scramble control (i-iv) and *IER3IP1* KD (v-viii) cells stained with Phalloidin, Tubulin, Rab11 and DAPI. Scale bar=5µm

**Figure S6:**
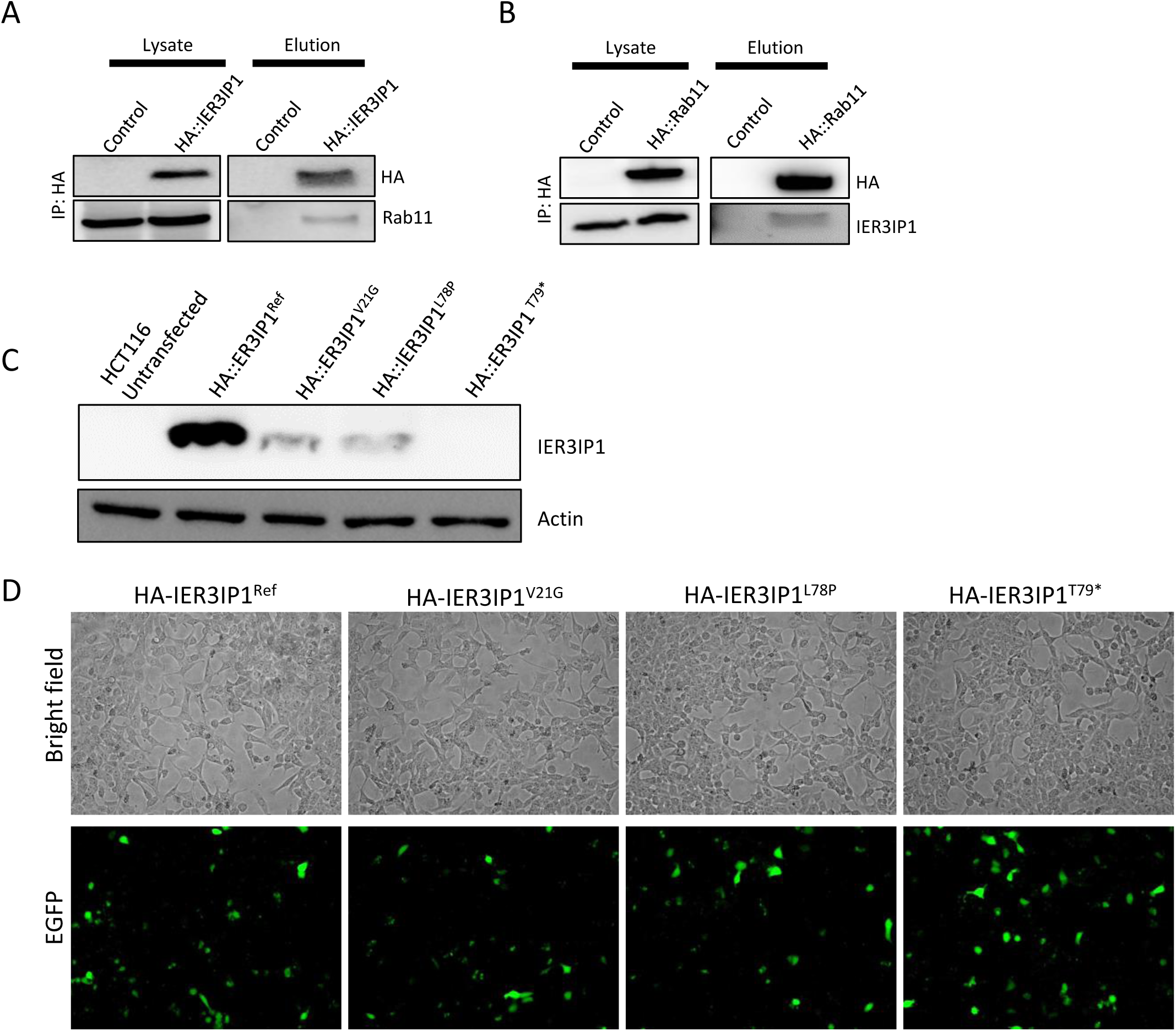
A. Immunoprecipitation assay showing Rab11 was immunoprecipitated from lysates of HEK293 cells expressing HA::IER3IP1 using anti-HA beads. Blots were probed with anti-HA and anti-Rab11 antibodies. B. Immunoprecipitation assay showing IER3IP1 was immunoprecipitated from lysates of HEK293 cells expressing HA::Rab11 using anti-HA beads. Blots were probed with anti-IER3IP1 and anti-HA antibodies C. Lysates from cells expressing *IER3IP1 Ref cDNA* and disease associated variants *V21G, L78P* and *T79\** were probed with anti-IER3IP1 and anti-actin antibodies D. HCT116 cells co-transfected with plasmid expressing *IER3IP1 Ref cDNA* and disease associated variants *V21G, L78P* and *T79\** and pADTrack-EGFP to check the transfection efficiency. Note that the number of GFP positive cells is almost the same for all the constructs suggesting that the difference in protein levels is not due to the differences in transfection efficiency.

